# Cell-Type Specific Interrogation of CeA *Drd2* Neurons to Identify Targets for Pharmacological Modulation of Fear Extinction

**DOI:** 10.1101/224261

**Authors:** Kenneth M. McCullough, Nikolaos P. Daskalakis, Georgette Gafford, Filomene G. Morrison, Kerry J. Ressler

## Abstract

Behavioral and molecular characterization of cell-type specific populations governing fear learning and behavior is a promising avenue for the rational identification of potential therapeutics for fear-related disorders. Examining cell-type specific changes in neuronal translation following fear learning allows for targeted pharmacological intervention during fear extinction learning, mirroring possible treatment strategies in humans. Here we identify the central amygdala (CeA) *Drd2*-expressing population as a novel fear-supporting neuronal population that is molecularly distinct from other, previously identified, fear-supporting CeA populations. Sequencing of actively translating transcripts of *Drd2* neurons using translating ribosome affinity purification (TRAP) technology identifies mRNAs that are differentially regulated following fear learning. Differentially expressed transcripts with potentially targetable gene products include *Npy5r, Rxrg, Adora2a, Sst5r, Fgf3, Erbb4, Fkbp14, Dlk1*, and *Ssh3*. Direct pharmacological manipulation of NPY5R, RXR, and ADORA2A confirms the importance of this cell population and these cell-type specific receptors in fear behavior. Furthermore, these findings validate the use of functionally identified specific cell populations to predict novel pharmacological targets for the modulation of emotional learning.

## Introduction

The amygdala is a mediator of the acquisition and expression of learned associative fear^1,2^. Composed primarily of GABAergic medium spiny neurons, the central amygdala (CeA) is intimately involved in controlling the expression of fear related behaviors^3,4^. Each of the CeA’s three main sub-nuclei (lateral capsular (CeC), lateral (CeL),and medial (CeM)) play distinct roles in specific behaviors and contain molecularly distinct sub-populations that have further behavioral specializations^5–10^. In the present set of experiments, we utilized Pavlovian fear conditioning, a paradigm used extensively for studying associative fear memories formed by the pairings of conditioned stimuli (CS; e.g. a tone) and unconditioned stimuli (US; e.g. a mild foot shock)^11–13^. Learned fearful associations may be ‘extinguished’ with additional unreinforced presentations of the CS alone, a process that closely resembles the clinical practice of exposure therapy used in treating individuals with PTSD. A promising area of treatment in PTSD includes the pharmacological enhancement of exposure-based therapies^14^. The aim of this study is to harness cell-type specific molecular techniques in order to identify more specific and effective pharmacotherapies for the treatment of fear-related disorders.

Foundational research as well as more recent analyses highlight the striatum-like nature of the central amygdala^15^. Striatal dopamine receptor 1 (*Drd1*) populations (direct pathway neurons) promote movement, while dopamine receptor 2 (*Drd2*) populations (indirect pathway neurons) inhibit movement^16,17^. Within the posterior CeA, it has been reported that corticotropin releasing factor (Crh), tachykinin 2 (*Tac2*), somatostatin (Sst), and neurotensin (Nts) expressing populations are contained within the larger *Drd1* expressing neuron population that promote directed motivational behaviors under certain conditions ^18,19^. Conversely, within the anterior CeC, the protein kinase C-δ (*Prkcd*) and calcitonin receptor-like (*Calcrl*) co-expressing population has been reported to be a sub-population of *Drd2* neurons mediating defensive behaviors or inhibiting motivated behaviors^19,20^. Given its potential role in fear behavior, the CeA *Drd2* expressing population is a high value target for translational investigation.

The dopaminergic system is well known for its role in appetitive learning; however, more recently it has been recognized for its importance in fear acquisition and fear extinction learning been equivocal on the^21–23^. Perturbations in the dopaminergic system have been implicated in the disease etiologies of several human pathologies ranging from Parkinson’s disease to schizophrenia, depression and posttraumatic stress disorder (PTSD)^24–26^. Although the dopamine receptor 2 (D2R) is clearly involved in fear acquisition and fear extinction learning, the literature to date has been equivocal on the role of D2R in the CeA, as different study designs demonstrate D2R antagonist administration may lead to conflicting effects^27–29^. In the present study, we separate the role of CeA *Drd2*-expressing neurons in fear behavior from that of receptor activity of D2R itself, and in doing so, indentify a large number of alternative gene targets that are modulated by fear learning.

The present study takes the most translationally direct approach by behaviorally and molecularly characterizing the CeA *Drd2* neuronal population, examining translational changes in this population following a fear learning event, and then pharmacologically manipulating identified targets at a clinically relevant time-point, during fear extinction. Molecular characterization of this population clearly identifies it as a unique population that is largely non-overlapping with other,previously described CeA populations. Direct chemogenetic enhancement of excitability in CeA *Drd2* neurons resulted in significantly enhanced fear expression. Translating ribosome affinity purification (TRAP) and sequencing of actively translated RNAs in the *Drd2* neuron population following fear conditioning yielded a diverse set of genes differentially regulated by behavior^30^. These differentially regulated genes included *Adora2a, Rxrg, Sst5r, Npy5r, Fgf3, Erbb4, Gpr6, Fkbp14, Dlk1* and *Ssh3.* Using the Druggable Genome database, genes with known pharmacological interaction partners were chosen and pharmacologically manipulated at a clinically relevant time point to oppose fear conditioning dependent changes, during fear extinction. Consistent with the identification of the *Drd2* expressing population as a fear expression supporting population, blockade of A_2A_R (G_αs_) or NPY5R (G_αi_) during fear extinction suppressed and enhanced fear expression respectively. Additionally, activation of RXR enhanced fear extinction consolidation. Together these data provide promising new targets for understanding and manipulating fear processes, and also demonstrate the power of identifying novel pharmacological targets through the use of cell-type specific approaches to amygdala circuit function.

## Results

### *Drd2* defines a distinct CeA population

Many molecularly distinct subpopulations have been identified across the CeA. Using RNAScope technology, we performed fluorescence *in situ* hybridization (FISH) in order to examine the *Drd2* population in relation dikkopf-related protein 3(Dkk3), dopamine receptor 1a (*Drd1a*), adenosine A2A receptor (*Adora2a*), corticotropin releasing factor (*Crh*), neurotensin (*Nts*), protein kinase C-δ (*Prkcd*), somatostatin (Sst), and tachykinin 2 (*Tac2).* Within the striatum, *Drd1a* and *Drd2* have expected intermingled, non-overlapping, expression patterns (Figure 1A-F). *Dkk3* strongly labels a population of BLA primary neurons. *Drd1a* strongly labels intercalated cell masses (ITC), especially the main intercalated island, and weakly labels some BLA cells. *Drd2* does not label any BLA or ITC cells. Within the CeA at anterior positions,*Drd2* primarily labels populations in the CeC and CeL with lower expression within the CeM, while *Drd1* primarily labels populations in the CeL and CeM with less expression within the CeC ^2,31,32^. At higher magnification it is clear that within the CeA *Drd1a* and *Drd2* maintain their non-overlapping expression with very few identifiable co-expressing cells (Figure 1G-L). *Drd2* is known to strongly co-express with *Adora2a* in the striatum. Similarly, we find an almost complete co-expression of *Drd2* and *Adora2a* within amygdala neurons.

**Figure 1.**
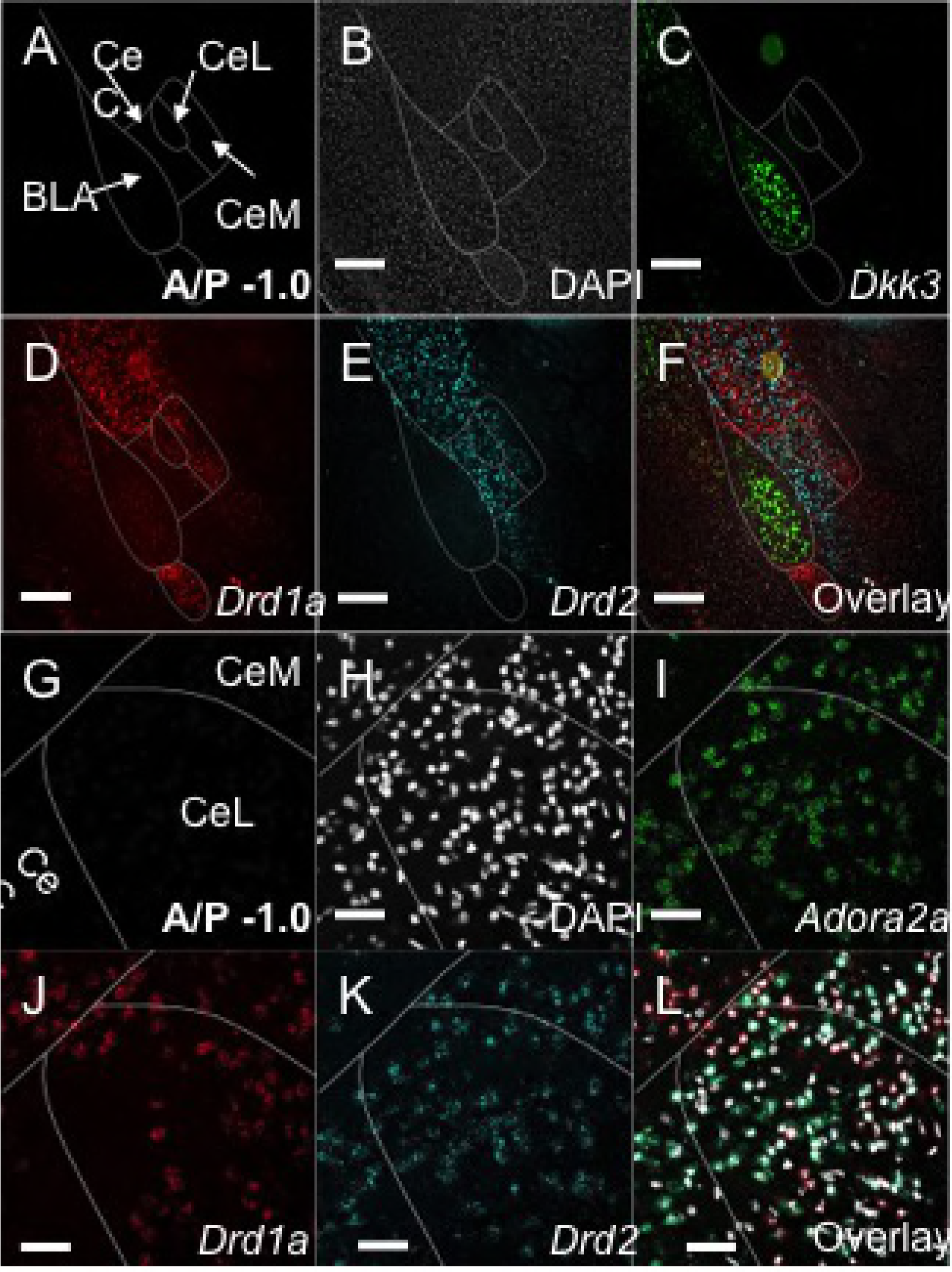
Comparison of CeA *Drd2, Drd1a*, and *Adora2a* populations. Expression of *Dkk3, Drd2, Drd1a*, and *Adora2a* were examined with FISH (RNA Scope, ACD Biosystems). **A**. Schematic of amygdala compartments within the temporal lobe. **B**. DAPI (Grey). **C**. *Dkk3* (Green) is expressed in a population of BLA pyramidal neurons. D. *Drd1a* (Red) is expressed in striatum, weakly in some BLAcells, ITC’s (especially Im), strongly in CeL and CeM, but weakly in the CeC. **E**. *Drd2* (Cyan) is expressed in striatum, CeC, CeL, but weakly in the CeM and not ITCs or BLA. **F**. Overlayof B-E. The dorsal CeA especially CeL expresses both *Drd2* and *Drd1a* populations; however theses populations segregate primarily to the CeC and CeM , respectively, more ventrally **G**. Schematic of higher magnification anterior dorsal region of CeA. **H**. DAPI (Grey). **I**. *Adora2a* (Green) is expressed strongly CeC, CeL and dorsal CeM. **J**. *Drd1a* (Red) is expressed strongly in CeL and CeM, but little expression is found in CeC. **K**. *Drd2* (Cyan) is expressed strongly in CeC, CeL and dorsal CeM. **L**. Overlay of H-K. *Adora2a* and *Drd2* entirely co-express. Very few examples of co-expression between *Drd1a* and either *Drd2* or *Adora2a* are found. Scale Bar: A-F 500 um, G-L 50 uM.

The anterior to posterior (A/P) position within the CeA has emerged as a strong potential variable when examining the behavioral functions of CeA neurons^19^. Therefore, the distribution of *Drd2, Drd1a*, and *Adora2a* expressing cells was examined across the length of the CeA (Supplemental Figure 1 A-X). Consistently, *Drd2* and *Drd1a* label non-overlapping populations, while *Drd2* and *Adora2a* label almost entirely overlapping populations with some single labeled cells found at the far ventral portion of the CeC. At anterior positions (A/P:-.82 to -1.2), *Drd2* strongly labels large populations within the CeC and CeL and to a lesser extent the CeM (Supplemental Figure 1A). Likewise, *Drd1a* labels populations within the CeL and CeM and many fewer cells in the CeC. At more posterior positions (A/P: -1.3 to -1.6) labeled cell distributions are less defined; the CeC, CeL and CeM are sparsely labeled aside from a strongly labeled dorsal *Drd2/Adora2a* population that appears to be contiguous with the striatum. Interestingly, the posterior CeA, especially posterior CeL, which contains the densest labeling for *Crh, Nts, Sst, Prkcd*, and *Tac2*, is only sparsely labeled with *Drd1a* (Supplemental Figure 1. D, H, L, P, T, X and Figure 2 M, N, Q, R).

To statically assess the extent to which the *Drd2* population overlaps with markers of other identified fear-related CeA populations, co-expression of *Drd2* with *Crh, Nts, Sst, Prkcd*, and *Tac2* was quantitatively assessed across the A/P axis of the CeA (Figure 2 A-Z and Supplemental Figure 2 A-J). Anterior CeA was considered to be between -.8 and -1.2 A/P while posterior CeA was considered as between -1.3 and -1.6. Posterior to approximately -1.6 was not examined, as the CeM is absent. Positive expression within a cell was visually scored as having five or more labeled puncta within twice the diameter of the nucleus. Single labeled images were scored then identified nuclei were overlaid and counted for none, single and double labeling.

**Figure 2.**
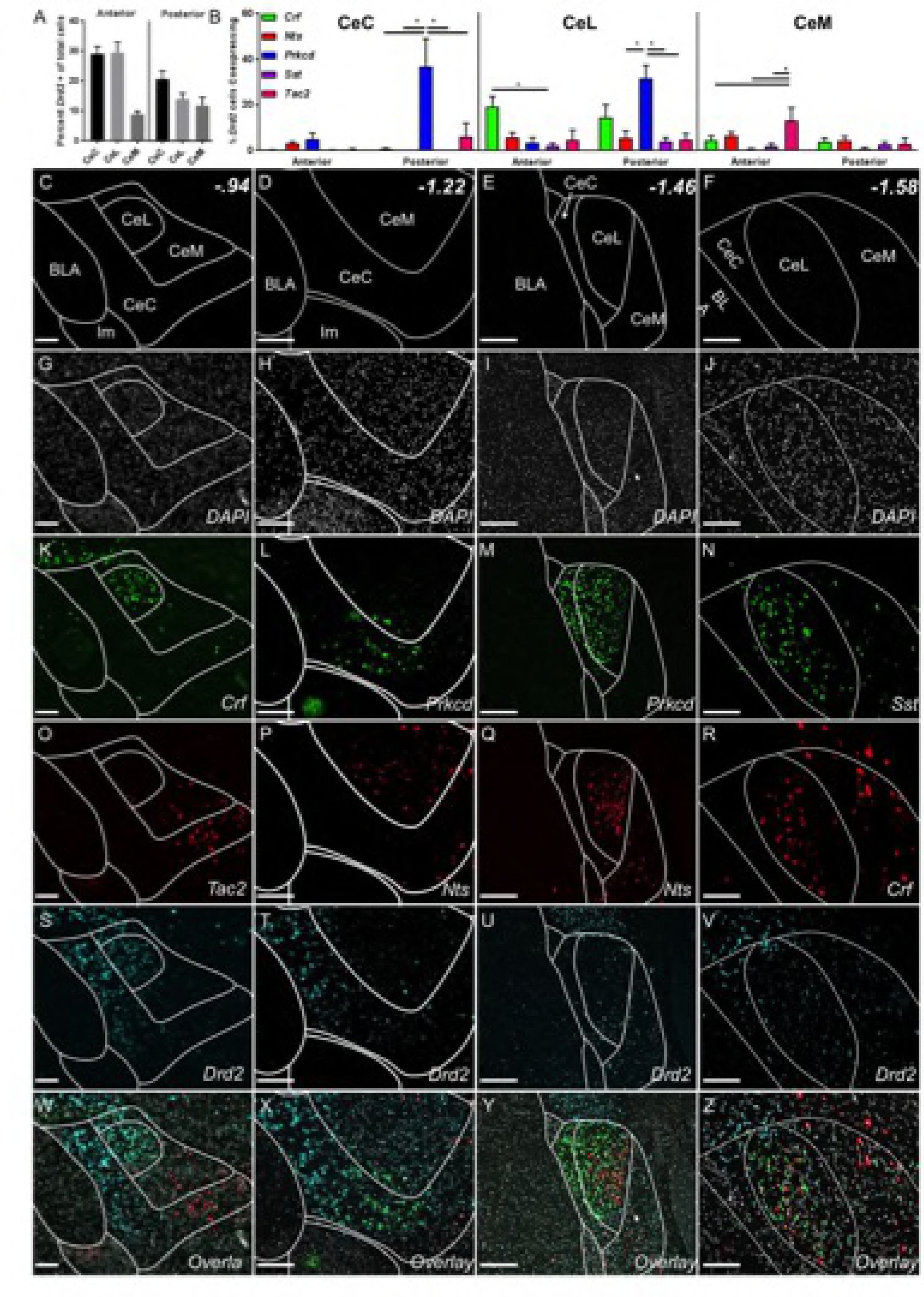
Co-localization of Drd2 with *Crh, Nts, Prkcd, Sst*, and *Tac2*. *Drd2* does not strongly co-express with any markers examined in anterior CeA. *Drd2* moderately co-expresses with *Prkcd* in posterior CeC and CeL. **A**. Density of *Drd2* cell population across anterior and posterior CeA represented as a percentage of total DAPI labeled cells. The strongest *Drd2* expression is found in anterior CeC and CeL. **B**. Quantification of *Drd2* co-expression with different CeA markers at anterior and posterior positions as a percentage of total *Drd2* expressing cells (CeC: 2-way ANOVA with anterior vs. posterior set as row factor (F(1,79)=13.2), p=.0005) and individualRNAs set as column factor (F(4,79)=16.19, p<.0001). Interaction (F(4,79)=10.56, p<.0001) and Sidaks multiple comparisons test within row: posterior: Crh vs. Prkcd p < .0001, Nts vs. Prkcd p < .0001, Sst vs. Prkcd p< .0001, and Tac2 vs. Prkcd p < .0001; CeL 2-way ANOVA with anterior vs. posterior set as row factor (F(1,74)=2.817), p=.0975) and individual RNAs set as column factor (F(4,74)=5.288, p<.0008). Interaction (F(4,74)=3.901, p<.0063)and Sidaks multiple comparisons test within row: anterior: Crh vs. Sst p < .05, posterior: Nts vs. Prkcd p< .005, Sst vs. Prkcd p < .0005, Tac2 vs. Prkcd p < .05; CeM: 2-way ANOVA with anterior vs. posterior set as row factor (F(1,77)=3.024), p = .086) and individual RNAs set as column factor (F(4,77)=2.578, p<.05). Interaction (F(4,77)=1.456, p = .22) and Sidaks multiple comparisons test within row: Posterior: Crh vs. Tac2 p < .04, Tac2 vs. Prkcd p < .005, Tac2 vs Sst p < .01). **C-F**. Map of CeA at A/P: -.94, -1.22, -1.46, and -1.58 respectively. **G-J**. DAPI (Grey) expression in at A/P: -.94, -1.22, -1.46, and -1.58 respectively. **K**. *Crh* (Green) is found primarily in CeL at A/P: -.84. **L**. *Prkcd* (Green) is found in a isolated population at the ventral aspect of the CeC at A/P -1.22. **M**. *Prkcd* (Green) is densely expressed in CeC and CeL at A/P: -1.46. **N**. *Sst* (Green) is densely expressed in CeL and more diffusely in CeM at A/P: -1.58. **O**. (Red) *Tac2* is expressed in ventral CeC and CeM at A/P: -.94. P. *Nts* (Red) is expressed almost exclusively in CeM at A/P: -1.22. **Q**. *Nts* (Red)is expressed densely in CeL and diffusely in CeM at A/P: -1.46. **R**. *Crh* (Red) is expressed densely in CeL and more diffusely in CeM at A/P: -1.58. **S**. *Drd2* (Cyan) is expressed strongly in CeC and CeL and more weakly in CeM at A/P: -.82. **T**. *Drd2* (Cyan) is expressed strongly in CeC and CeL and more weakly in CeM at A/P: -.94. **U-V**. *Drd2* (Cyan) is expressed more diffusely throughout CeA at A/P -1.46 and -1.58 respectively. **W-Z**. Overlay of G-V. Scale Bar: A-Z 100um.

Within the anterior CeA, *Drd2* was not found to extensively co-express with any other population examined (Figure 2B). Within the anterior CeL and CeM *Drd2* co-expressed significantly more with *Crh* and *Tac2* respectively compared to other markers, although total co-expression was low at 19.3% and 13.1% of *Drd2* positive cells co-localized with *Crh* and *Tac2*, respectively (Figure 2 C, G, K, O, S, W and Supplemental Figures 2C and G). Within the posterior CeC and CeL, *Drd2* co-expressed more with *Prkcd* than any other marker (36.7% and 31.5% of *Drd2* cells in the CeC and CeL, respectively); however this represented a relatively low percentage of total *Prkcd* positive cells (13.9% and 10.1% of *Prkcd* positive cells in the CeC and CeL, respectively)(Supplemental Figure 2B). Staining for *Prkcd* was found beginning in the anterior ventral CeC forming a contiguous population to a more dorsal position posteriorly where the traditionally reported CeC and CeL population fear extinction, we directly is found (Figure 2L & M).

### Chemogenetic activation of CeA *Drd2* neurons enhances fear expression

To determine the precise role of the *Drd2*-expressing population in fear extinction, we directly manipulated these neurons during extinction using designer receptors exclusively activated by designer drugs (DREADDs)^33^. Drd2-Cre (B6.FVB(Cg)-Tg(Drd2-Cre)ER43Gsat/Mmucd) mice and non-Cre expressing littermate controls were infected bilaterally with a Cre-dependent Gs-coupled DREADD virus (Figure 3A-C). Gs-DREADD expression was visualized through its mCherry tag (Figure 3B and C). Three weeks following infection, mice were mildly fear conditioned with 5 CS/US (0.4 mA US footshock) pairings to avoid ceiling effects (Figure 3D). A trend towards increased freezing in the Drd2-Cre mice was found during conditioning; if this represents a true finding it may have been caused by leakage of the Gs-DREADD; however, freezing during the final CS/US paring was very similar between both groups, suggesting no differences in overall fear learning. 30 minutes prior to the extinction session (15 CS), all mice were injected with CNO (1 mg/kg, i.p. in saline). Mice that expressed Cre-recombinase and thus expressed the Gs-DREADD in *Drd2* neurons exhibited significantly more freezing to the tone throughout the extinction session. Importantly, 24 h later, after a wash-out period when DREADDs were no longer active (previous research has shown that wash-out is 6-10 h^34,35,36^), Gs-DREADD expressing mice again displayed significantly more freezing to the CS compared to controls during a 30 CS extinction retention session. The rate of extinction of both groups in the initial extinction session did not significantly differ, suggesting that the enhancement in freezing during the second extinction session was likely due to blockade of extinction consolidation.

**Figure 3.**
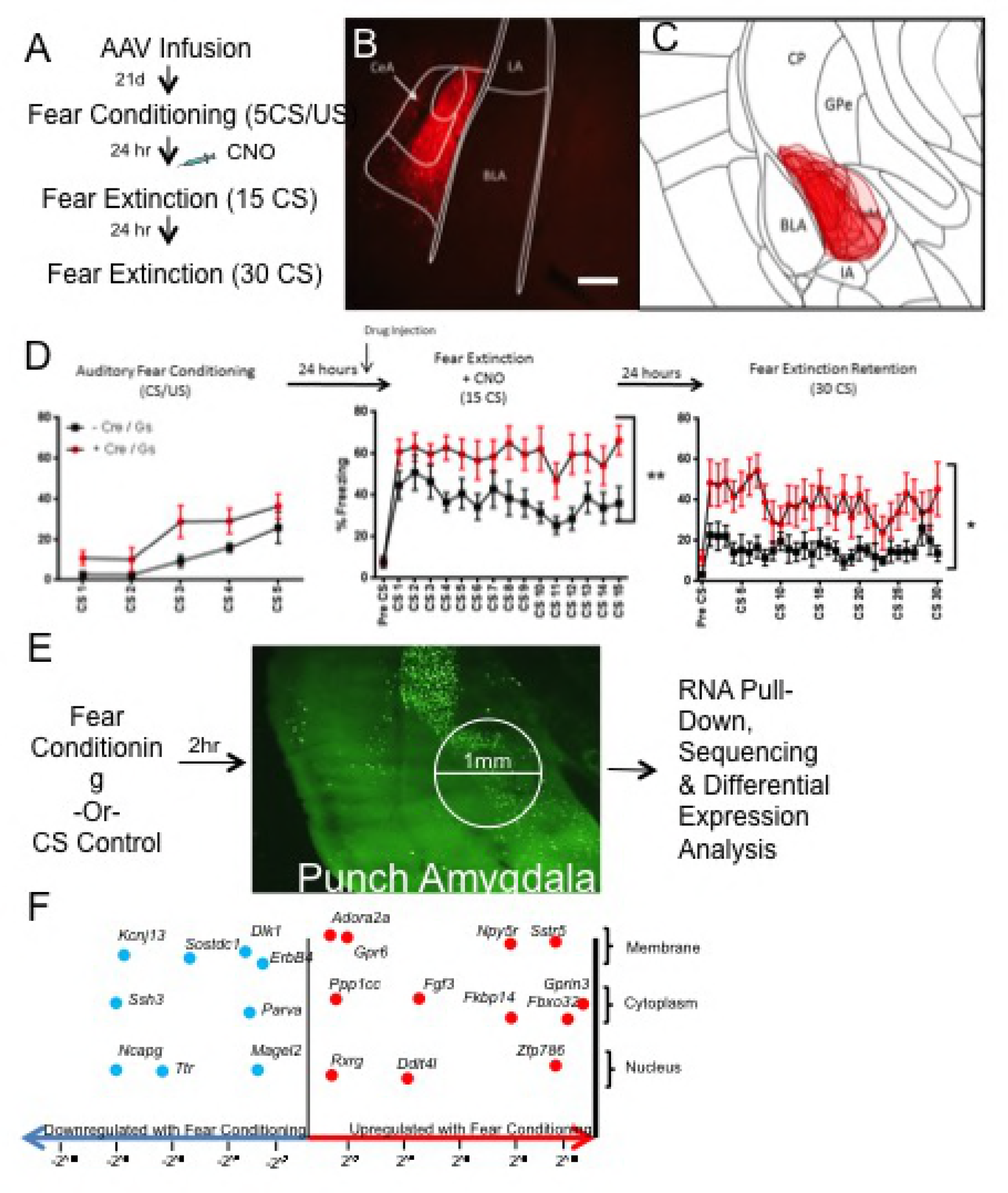
Cell-type specific manipulation of and TRAP isolation from CeA *Drd2* population. **A**. Schematic ofexperimental design. **B**. Representative expression pattern of mCherry-tag expression in *Drd2* neurons of theamygdala. Scale Bar: 200 um. **C**. Collapsed over-lay of expression pattern of mCherry for Cre-expressing experimental animals. Expression is generally constrained to CeC and CeL with limited expression in CeM. **D**. Mice were weakly fearconditioned to 5 CS/US pairings(6 kHz tone, .4 mA foo-shock) (2-way RM ANOVA F(1,18)= 4.382, p > .05). Mice were injected i.p. with CNO 30min prior to fear extinction session. Mice expressing Gs-DREADD-mCherry expressed significantly more fear during the entireextinction session than non-carrier controls (2-way RM ANOVA F(1,18)=11.49, ** p < .01). Twenty-four hours later during the second extinction (retention) session mice expressing Gs-DREADD-mCherry expressed significantly more fear (2-way RM ANOVA F(1,18)=7.512, * p < .05). **E**. Schematic of TRAP experiment. Animals were fear conditioned (5 CS/US, .65 mAfoot-shock) or exposed to training environment. 2 hours lateranimals weresacrificed, 1 mm punches centered over CeA were taken and TRAP procedure was completed. **F**. Selected differential expressionresults of fear conditioned vs. control animalswith log-fold change on x-axis. Genes found to be down-regulatedfollowing fear conditioning compared to controls (Blue, Leftward) include *Erbb4, Dlk1, Parva, Ssh3, Ttr*, and *Kcnj13.* Genes found to be up-regulated following fear conditioning compared to controls (Red, Rightward) include *Adora2a, Gpr6, Ppp1cc, Rxrg, Fgf3, Npy5r, Sstr5, Fkbp14* and *Gprin3.*

### Characterization of dynamic mRNA changes in *Drd2* cells following fear conditioning

To further characterize the Drd2-expressing population, we next examined expression changes in *Drd2* neurons following fear conditioning. Transcripts were examined following fear conditioning based on the expectation of this time point predicting the direction of protein expression levels 24-hours later prior to fear extinction. Additionally, we expected that modulation of fear learning precipitated molecular changes may lead to decreased fear expression or enhanced extinction. To identify actively translating mRNA transcripts, TRAP protocol was utilized^30^. The Drd2-Cre mouse line was crossed with the floxed-stop-TRAP (B6.129S4-Gt(ROSA)26Sortm1(CAG-EGFP/Rpl10a,-birA)Wtp/J) line to generate a double transgenic line, Drd2-TRAP. Expression of the L10a-GFP transgene closely recapitulated our observed expression patterns of *Drd2* (Supplemental Figure 3)^37–39^. Next, animals were either fear conditioned (5 CS/US tone-shock pairings with 0.5 sec, 0.65 mA foot shock US) or exposed to the tone CS in the chamber in the absence of any US. Fear conditioned animals exhibited expected increases in freezing responses to the CS (Supplemental Figure 4A). Animals were then sacrificed 2 h following behavior, micropunches centered over the CeA were collected, and TRAP was performed to obtain isolated mRNA from *Drd2* neurons (Figure 3E). High quality RNA was retrieved from the TRAP protocol (RIN=8.5-10). To verify the specificity of RNA pull-down, qPCR analysis of samples was performed to compare bound versus unbound samples. Ribosomal subunit 18S was found at higher levels in the bound fraction compared to the unbound fraction, confirming enrichment for ribosomes (Supplemental Figure 5A). When expression levels of *Drd2* and *Drd1a* were compared in ribosomal bound and unbound fractions, the bound fraction had a large enrichment of *Drd2* versus *Drd1a* transcripts when compared to the unbound fraction (Supplemental Figure 5B) ^40–42^. Ribosomes specifically expressed in *Drd2* neurons were successfully pulled down and RNA collected from these pull-downs demonstrated expected characteristics of *Drd2* neurons; strong expression of *Drd2* and weak expression of *Drd1a*.

Sequencing of RNA collected from *Drd2* neuron ribosomes revealed genes dynamically regulated following fear conditioning, many of which have been previously reported to be involved in fear and anxiety-like behaviors (Figure 3F). False discovery rate (FDR) adjusted p-values were calculated and FDR of 5% and fold-change of 2^^0.5^ cut-offs were set(Full list of differentially expressed genes is in Supplemental Table 1). Using, the Mouse Gene Atlas dataset, initial analysis using Enrichr confirms amygdala specificity of pull-down and gene change (Supplemental Table 2)^43^. Further enrichment analysis using Jensen Compartments dataset confirms neuronal specificity of pull down and gene change (Supplemental Table 3). Consistent with activity dependent gene changes, MetaCore Gene Ontology Processes identifies neuronal developmental and adenylate cyclase related processes as highly significantly recruited by fear conditioning (Figure 4A). MetaCore Gene Ontology Diseases identifies Schizophrenia and nervous system diseases as gene categories most related to gene changes in *Drd2* neurons (Figure 4B). Interestingly, GSEA identified gene group differences in the entire RNA-seq dataset as most concordantly similar, but in the opposite direction to and two gene data sets identified in hippocampus and mPFC of humanized 22q11.2 deletion model of Schizophrenia (Supplemental Figure 6)^44,45^. This is informative for interpreting the enrichment of our top FDR-significant genes with Schizophrenia disease set by MetaCore. Weighted network analysis was completed to examine differential expression of *Drd2* genes in the context of human PTSD.Using GeneMANIA Cytoscape, differentially expressed transcripts were mapped into a self-organizing weighted network, where all of the genes were interlinked at multiple levels (co-expression, physical interactions, common pathway) (Figure 4C)^46,47^. Over-all gene network analysis reveals that differentially regulated genes are primarily co-expressed and are part of common ontologies without belonging to a single dominant pathway. To identify potential targets for pharmacological manipulation, differentially expressed genes were examined for the availability of agonists or antagonists using MetaCore Drugs for Drug targets tool (Supplemental Table 4). Finally, potential drug targets were examined in the literature for being high quality, blood-brain barrier penetrant, agonists or antagonists. Using this identification approach, inclusive of our *a priori* interest in *Adora2a* among other potential *Drd2*-neuron specific genes (above studies), *Npy5r*, and *Rxrg* were selected for further pharmacological examination^48–51^. Additional markers found to be modulated with fear learning that may be of further interest also include *Sst5r, Fgf3, Erbb4, Gpr6, Fkbp14, Dlk1* and *Ssh3.*

**Figure 4.**
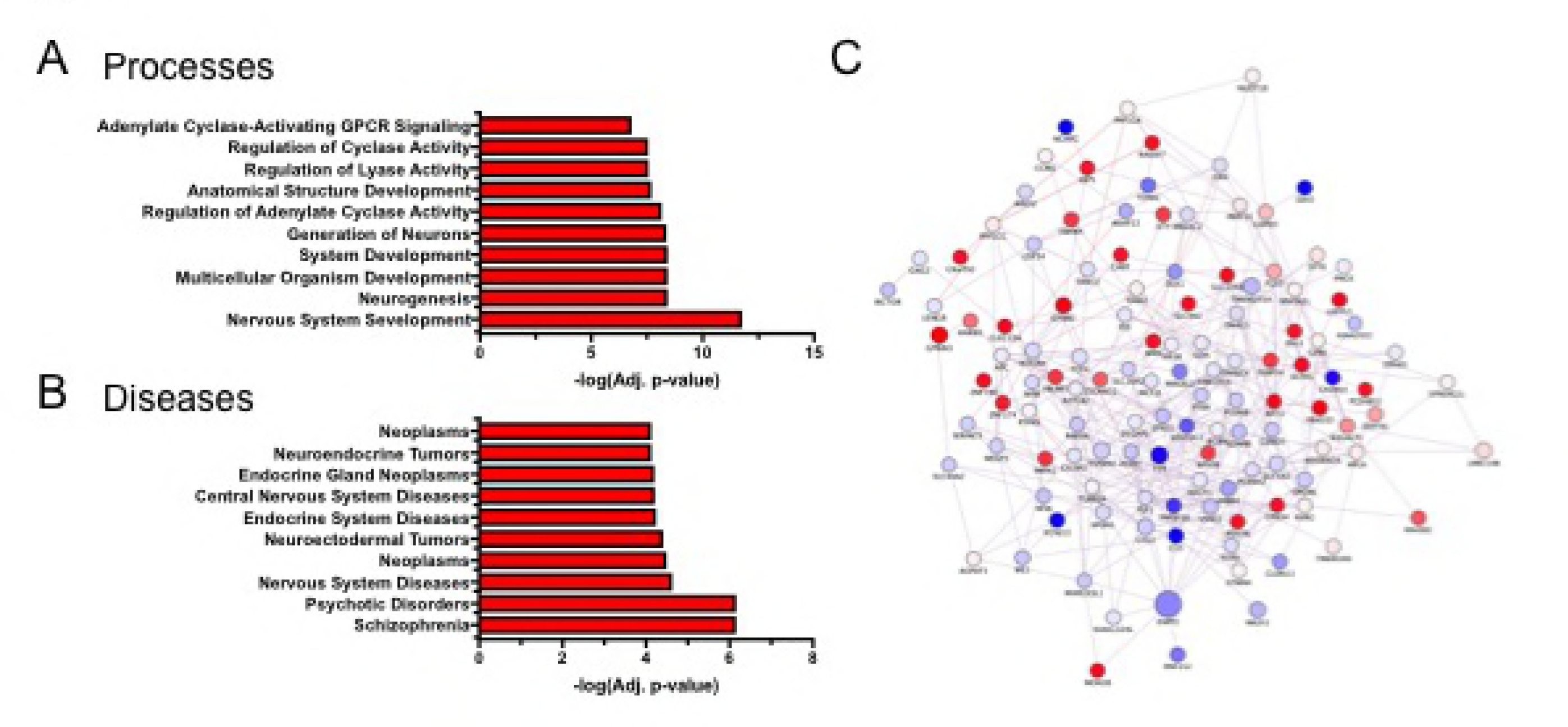
Bioinformatic analysis of differentially expressed genes in *Drd2* population following fearconditioning. **A**. Enrichment analysis for the MetaCore Gene Ontology Processes identifies highly significant processes related to gene changes in *Drd2* neurons. **B**. Enrichment analysis for the MetaCore Gene Ontology Diseases identifies highly significant diseases related to gene changes in *Drd2* neurons. **C**. Weighted Network ofgenetically annotated transcripts showing differential gene expression. Differentially expressed transcripts were analyzed with the GeneMania Cytoscape plug-in using the default setting, but without extending the network with additional nodes. Genes that were not connected with others are not represented. The node size represents the-log(FDR-adjusted P-value), whilethe intensity of the color represents logFC (red nodes denote up-regulation in FC, while blue nodes denote down-regulation). The between-nodes edges represent relationships, the color of the edges represent the type of the relationship (76.5% coexpression in purple, 22% physical interactions in pink, 1.5% common pathway interactions in light blue), and the thickness of the edges denotes weight (i.e., strength of the pairwise relationship).

Based upon the altered translational activity in *Drd2* neurons following fear conditioning, one potential route to enhance fear extinction is to pharmacologically manipulate the activity of the identified translated protein products. ADORA2A, NPY5R, and RXR were chosen as potential targets for pharmacological modulation of fear extinction, as they were robustly differentially expressed in the *Drd2* fear-regulating neuronal population, and they have well-understood mechanisms of action, making them attractive targets for pharmacological manipulation of fear extinction.

### Manipulation of ADORA2A, NPY5R, and RXR recapitulates the role of *Drd2* neurons in fear behavior

Agonists and antagonists targeting ADORA2A, NPY5R, and RXR receptors were chosen from the literature (Figure 5A)^52^. *Adora2a* was an attractive candidate for further inquiry and was chosen for initial characterization based on a number of reasons; 1) it has previously been shown to almost entirely co-express with *Drd2* (Figures 1G-L and 2) within the amygdala, and 2) several pharmacological agents targeting ADORA2A are currently in clinical trials or have been approved for use in humans^53,54^. The highly selective ADORA2A antagonist, istradefylline, is selective for ADORA2A over ADORA1 with a Ki of 2.2 and 150 nM respectively^55,56^.

**Figure 5.**
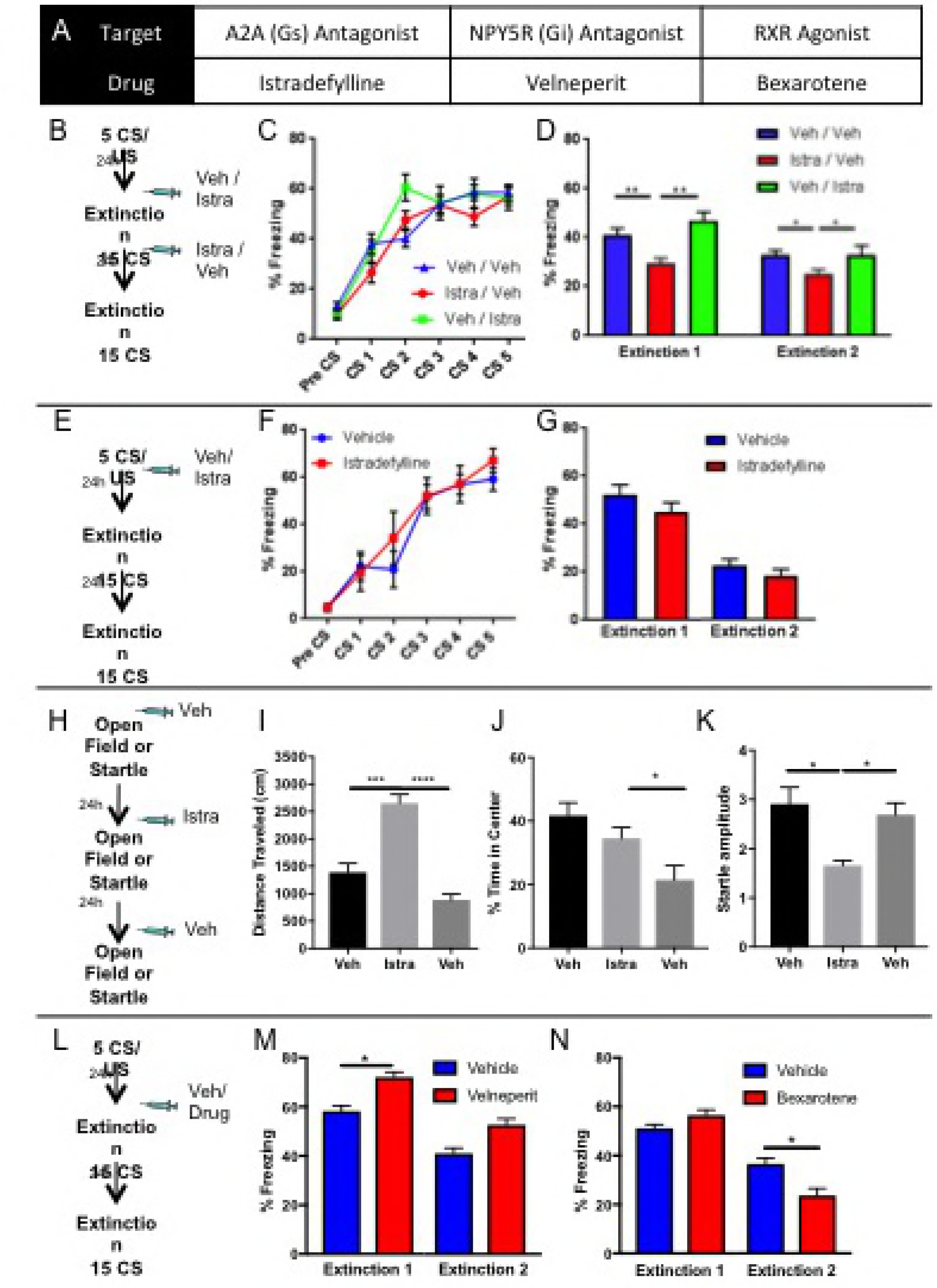
Pharmacological manipulation of ADORA2A, NPY5R, and RXR during behavior. *Adora2a, Npy5r*, and *Rxrg* were found to be increased following fear conditioning; therefore, the effect of pharmacological manipulation of ADORA2A, NPY5R, and RXR during fear extinction was examined to assess their utility as potential enhancers of exposure therapy. **A**. List of pharmacological agents used, their targets and the effects of binding to target. **B**. Schematic of experimental design for examination of ADORA2A antagonism by Istradefylline prior to or following fear extinction. **C**. Three groups of animals were fear conditioned (5 CS/US, .65 mA foot-shock). **D**. Pre-extinction injection of istradefylline (Istra/Veh group) causes significant decrease in freezing compared to vehicle injected controls (2-way RM ANOVA F(2,71)=10.26, p < .0001; Tukey’s Multiple Comparisons: I/V vs. V/I p = .0005, I/V vs. V/V p = .0017, V/I vs. V/V p = .517). Animals that previously received istradefylline prior to fear extinction (Istra/Veh) continue to express less freezing24-hours later during second extinction session (retention) compared to vehicle controls (Veh/Veh) and those that received istradefylline following extinction (Veh/Istra) (2-way RM ANOVA F(2,69)= 5.381 (2 animals removed b/c injuries fromfighting, 1 from V/V and 1 from I/V), p < .01; Tukey’s Multiple Comparisons: I/V vs. V/I p = .0236, I/V vs.V/V p = .0181, V/I vs. V/V p = .8988). **E**. Schematic of experimental design for examination of the effect ofADORA2A antagonism following fear conditioning. **F**. Two groups of mice were fear conditioned (5 CS/US, .65 mA foot-shock). G. No effect of prior istrafedylline treatment following fear conditioning was detected during first (2-way RM ANOVA F(1,10)=1.22, p>.05) or second extinction session (2-way RM ANOVA F(1,10)=0.88, p>.05). **H**. Schematic for experimental design of examination of effect of istradefylline on locomotion, center-time and acoustic startle. **I**. Pre-session administration of vehicle (Veh) or istradefylline (Istra). Day 2 istradefylline treatment caused acute increase in distance traveled compared to day 1 Vehicle that returned to baseline on day 3 Vehicle test (RM ANOVA F(1.443,8.656)=60.77 p < .0001; Tukey’s Multiple Comparisons: Veh (D1) vs Istra p=.0009, Veh (D1) vs. Veh (D3) p= .0984, Veh (D3) vs. Istra p < .0001). **J**. Pre-session administration of vehicle (Veh) or isradefylline (Istra). Day 2 istradefylline treatment did not cause changes in anxiety like behavior (time in center). Day 3 Veh did have reduced time incenter compared to Day 2 Istra; however, this is likely due to habituation (RMANOVA F(1.542, 9.253)=6.602 p < .05;Tukey’s Multiple Comparisons Test Veh (D1) vs Istra p=.4570, Veh (D1) vs. Veh (D3) p= .0577, Veh (D3) vs. Istra p=.0440). **K**. Pre-session administration of vehicle (Veh) or isradefylline (Istra). Day 2 treatment with istradefylline caused a decreased acoustic startle amplitude that did not persist into Day 3 vehicle treatment (RM ANOVAF(1.794,16.14)=8.203 p = .0043; Tukey’s Multiple Comparisons Test Veh (D1) vs Istra p=.0205,Veh (D1) vs. Veh (D3) p= .7924, Veh (D3) vs. Istra p=.0111)). **L**. Schematic for experimental design of examination of effects of venelperit and bexarotene. **M**. Cohorts of mice were fear conditioned (5 CS/US, .65 mA foot-shock), no differences between groups was detected. Pre-extinction injection of velneperit caused increased freezing when compared to control vehicle group, Extinction 1 (2-way RM ANOVA, F(1,13)=15.74, p < .005). No difference between groups was detected 24-hours later during second extinction session, Extinction 2. **N**. Cohorts of mice were fear conditioned (5 CS/US, .65 mA foot-shock), no differences between groups was detected. Pre-extinction injection of bexarotene causes no within sessionchange in behavior, Extinction 1. Twenty-four hours later during Extinction 2, animals that had previously been injected with bexarotene prior to Extinction 1 expressed significantly less freezing than vehicle injected controls (2-way RM ANOVA, (1,29)=5.761, p < .05).

To examine the effect of ADORA2A antagonism on fear extinction three cohorts of mice were fear conditioned(5 CS/US, 0.65 mA foot shock)(Figure 5B and C). Twenty-four hours following fear conditioning, and 30 minutes prior to fear extinction, mice were injected with istradefylline (3 mg/kg) (group I/V) or vehicle (10% DMSO, 1% NP-40 in saline i.p.) (groups V/V and V/I) (Figure 5B). Additionally, immediately following fear extinction (15 CS) mice were again injected with istradefylline (3 mg/kg) (group V/I) or vehicle (groups I/V and V/V). Injection of istradefylline but not vehicle prior to fear extinction (15 CS, Extinction 1) greatly decreased freezing during extinction training when drug was on-board (Figure 5D). Twenty-four hours later, following drug clearance, mice that had previously been injected with istradefylline prior to fear extinction, but not those injected following it, expressed significantly less freezing during a second extinction session (15CS, Extinction 2). These data suggest that blockade of ADORA2A during fear extinction, but not during extinction consolidation, is sufficient to enhance fear extinction learning.

To further examine the role of ADORA2A in fear consolidation, a separate cohort of mice was fear conditioned (5 CS/US, .65 mA) and given injections of istradefylline (3 mg/kg, i.p.) or vehicle directly following the fear conditioning training session (5 CS/US, 0.65 mA foot shock), (Figure 5 E and F). Although there was a trend towards decreased fear expression during a fear extinction (15 CS) session 24 hours later in mice treated with istradefylline, no significant effect of ADORA2A blockade on fear consolidation was detected.

Istradefylline is a potential drug treatment for Parkinson’s disease, thus it is possible that locomotor effects obscured the effects of drug on fear behavior; therefore, two cohorts of mice were tested for locomotor and anxiety-like behaviors in an open field, and acoustic startle responses on consecutive days (Figure 5H). Injection of istradefylline (3 mg/kg) but not vehicle significantly increased the distance traveled in the open field (Figure 5I); however 24 hours later the distance traveled had returned to pre-treatment levels. Importantly, increased distance traveled was not accompanied by any anxiogenic or anxiolytic effects in the open field. The decreased locomotor activity across days is likely due to habituation to the chamber context. Finally, istradefylline acutely decreased baseline acoustic startle; however this effect was not present 24 hours later when startle amplitude returned to pre-treatment levels. Together, these data suggest that the effects of istradefylline in enhancing extinction retention tested 24 hours after drug administration are unlikely the result of alterations in locomotion or effects on anxiety like behavior, per se.

NPY5R and RXR were additional identified targets that were examined for pharmacological enhancement of fear extinction. Velneperit antagonizes NPY5R, while Bexarotene is a RXR agonist (Figure 5L). A cohort of animals was fear conditioned (5 CS/US, 0.65 mA foot shock) (Supplemental Figure 7A). Twenty-four hours later, 90 minutes prior to fear extinction (15 CS), animals were given injections of Velneperit (100 mg/kg) or Vehicle (DMSO) (Figure 5M). Animals treated with Velneperit (NPY5R antagonist) expressed significantly more freezing than animals injected with vehicle (Extinction 1). Twenty-four hours later during a second extinction session (Extinction 2), no difference between groups was detected. Another cohort of animals was fear conditioned (5 CS/US, 0.65 mA foot shock) (Supplemental Figure 7B), and 24 hours later, 90 minutes prior to fear extinction (15 CS, Extinction 1), animals were given injections of Bexarotene (RXR agonist, 50 mg/kg) or Vehicle (DMSO) (Figure 5N). No differences between groups were detected. Twenty-four hours later during a second extinction session (Extinction 2) animals previously treated with Bexarotene prior to Extinction 1 expressed significantly reduced fear compared with controls. These pharmacological agents predictably affected fear extinction learning in a manner consistent with our hypothesized role for the *Drd2* population being a fear supporting population whose activation or inhibition is sufficient to modulate fear. Antagonizing G_αs_-coupled ADORA2A dramatically decreased fear expression, as would be expected by decreasing activity of a fear-supporting population. In contrast, antagonizing G_αi_-coupled NPY5R increased fear expression as would be expected by decreasing inhibition of (increasing activity of) a fear-supporting population. Activation of RXR may act to generally enhance extinction consolidation as observed with Bexarotene treatment, although the mechanism by which this may occur is unclear as RXRs are nuclear hormone receptors with a variety of binding partners^57^.

### Dynamic regulation of *Drd2* after fear extinction

*Drd2* expression was not significantly changed after fear conditioning in the above reported TRAP study; however, the literature suggests that D2R is involved in the control and consolidation of fear and extinction learning. Therefore, the dynamic regulation of *Drd2* was examined after fear extinction. Two groups (FC 1 and FC 30) of animals were fear conditioned (5 CS/US, 0.65 mA foot shock)(Supplemental Figure 8A). Twenty-four hours later three groups received differing CS exposures; FC30 received 30 CS presentations; FC1 received 1 CS presentation and remained in the chamber for the remainder of the session; HC30 received exposure to 30 CS presentations with no previous training experience. A home cage (HC) control group was also included. Each cohort of mice was sacrificed 2 hours following extinction training, RNA was isolated from 1 mm micropunch centered over the CeA, and *Drd2* expression levels were examined via qPCR (Supplemental Figure 8B and C). *Drd2* mRNA expression was significantly increased in the extinction group (FC30) when compared to all other groups and no significant change from HC was found in either HC30 or FC1 groups. These data suggest that dynamic regulation of *Drd2* may be involved in the consolidation of fear extinction, potentially increasing inhibition of this population through G_αi_ coupled D2 receptors. This dynamic regulation with fear extinction consolidation is consistent with our findings of differential modulation of extinction learning with targeted Drd2-cell type specific pharmacological approaches.

## Discussion

The present study: 1) Examined the distribution and co-expression of *Drd2* with *Drd1a, Adora2a, Crh, Nts, Sst, Prkcd*, and *Tac2* across the A/P axis; 2) Identified *Drd2* expressing neurons as a fear supporting population through direct chemogenetic manipulation; 3) Characterized cell-type specific translational changes following fear conditioning and identified many dynamically regulated genes including *Adora2a, Npy5r, Rxrg, Sst5r, Fgf3, Erbb4, Fkbp14, Dlk1*, and *Ssh3*; and finally, 4) Pharmacologically manipulated ADORA2A, NPY5R, and RXR to assess their viability as potential *Drd2* cell-type specific targets for pharmacological enhancement of fear extinction.

The identification of a fear supporting population in the CeC is consistent with previous findings that the CeC specifically receives input from the fear promoting prelimbic cortex as well as other anxiety and pain related areas^58,59^. Our findings of strong co-expression of *Drd2* and *Adora2a* but not *Drd1a* are consistent with findings in other regions^42^. Interestingly, we found lower co-expression of *Drd2* with *Prkcd* in the posterior CeL compared to reports by De Bundel et al. and in the anterior CeC compared to reports by Kim et. al.^19,60^. The former instance is explained by De Bundel’s use of a Drd2::GFP reporter mouse; reporter mice may strongly express a transgene in cells that only express lower levels of the native transcript and thus were below our detection criteria. Likewise, discrepancies with Kim et al., are likely due to our use of stricter criteria for positive expression. In either case, data from both reports support our findings of *Drd2* as a fear supporting population. Another interesting discrepancy between our data and that reported by Kim et al., is that we demonstrate very little *Drd1a* expression in the posterior CeL. This is remarkable because the posterior CeL contains the densest *Crh, Nts, Sst, Prkcd*, and *Tac2*, populations that were reported to correspond with *Drd1a* neurons in this area. This discrepancy may again be due to our more strict criteria for positively expressing cells.

Overall, the presented behavioral data are remarkably consistent. Manipulation of the *Drd2* neuronal population either through Gs–DREADD, or the inhibition of ADORA2A (G_αs_) or NPY5R (G_αi_), drives fear expression in directions consistent with this being a fear supporting population. ADORA2a is known to be co-expressed with D2R and these receptors have been shown to have opposing actions, suggesting that both receptors may be viable candidates for modulation of a single sub-population^61,62^. An important consideration is that drugs were administered systemically, thus making it impossible to claim that effects were mediated exclusively through receptors found in the CeA. However, as the goal of this line of research is to identify potentially clinically relevant targets for enhancement of therapy, it is advantageous to test candidates as they would be used in the clinic, that is, systemically and prior to exposure therapy.

The finding of an acute increase in locomotion with global A_2A_R antagonism is consistent with reports in the literature and is expected because manipulation of the indirect pathway is a common treatment for Parkinson’s Disease^27,61^. Additionally, locomotor effects observed with the ADORA2A antagonist closely mirror results observed from direct DREADD manipulation of *Adora2a* neurons^63^. The transience of locomotor effects as well as the absence of effects on anxiety-like behavior suggest that changes in freezing during subsequent testing are due to effects on learning and are not the result of locomotor changes. These results are also consistent with reports that ADORA2A antagonism with SCH58261 produces deficits in contextual fear conditioning^64^.

Profiling changes in actively translating RNAs using TRAP protocols provides a unique window into the acute responses of these neurons to a learning event. We sought to identify transcripts that were differentially regulated following fear learning, so that changes in protein activity might be pharmacologically opposed at a later time point; during fear extinction.There are several other important time points to compare including prior to and following fear extinction, which will be important subjects for future investigation.

Bioinformatic analysis of TRAP-seq data emphatically confirms specificity of pull-down to amygdala neurons. Network analysis reveals that identified differentially expressed genes are primarily co-expressed. Although genes do not to a great extent belong to a single pathway, they are part of common ontologies suggesting domains of proteins that may be valuable to interrogate in the future. Several genes including *Adora2a, Sst5r, Npy5r, Fgf3* and *Erbb4*, have been directly implicated in or are in well-established signaling pathways implicated in the control of fear learning. Others genes such as *Rxrg, Gpr6, Fkbp14, Parva, Dlk1* and *Ssh3* have not been studied in the context of fear biology, but may provide valuable insights upon further investigation. Interestingly, several of these genes, most prominently *Adora2a* and *SstR5*, have been implicated in human anxiety disorders^65,66^.

Data presented here identifies potential pharmacological enhancers of extinction by leveraging cell-type specific techniques in a fear-controlling population. This approach represents a potential avenue for predicting novel targets for the modulation of emotional learning, generating more specific and effective treatments of psychiatric illnesses such as PTSD.

## Methods

### Animals

C57BL/6J mice were obtained from Jackson Laboratories (Bar Harbor, ME). B6.FVB(Cg)-Tg(Drd2-Cre)ER43Gsat/Mmucd mice were obtained from the MMRRC and produced as part of the GENSAT BAC Transgenic Project. Rosa26 fs-TRAP (B6.129S4-Gt(ROSA)26Sortm1(CAG-EGFP/Rpl10a,-birA)Wtp/J) mice were obtained from Jackson Laboratories. Drd2-TRAP mice were generated by crossing Drd2-Cre and Rosa26 fs-TRAP lines. All mice were adult (8-12 week) at the time of behavioral training. All mice were group housed and maintained on a 12hr:12hr light:dark cycle. Mice were housed in a temperature-controlled colony and given unrestricted access to food and water. All procedures conformed to National Institutes of Health guidelines and were approved by Emory University Institutional Animal Care and use Committee. Animal numbers were calculated using G*Power 3 software using previous experiments to inform expected means and standard deviations for expected large and medium effect sizes for chemogenetic and pharmacological manipulations respectively^1^. Animals were assigned to groups based upon genotype or randomized to treatment. Experimenter was blinded to genotype of animals. Blinding to drug administration was not possible; however, animal id’s were coded during data analysis.

### Surgical Procedures

Mice were deeply anesthetized with a Ketamine/Dexdormitor (medetomidine) mixture and their heads fixed into a stereotaxic instrument (Kopf Instruments). Stereotaxic coordinates were identified from Paxinos and Franklin (2004) and heads were leveled using lambda and bregma. For viral delivery (Figure 3), a 10μl microsyringe (Hamilton) was lowered to coordinates just above CeA (A/P -1.2, M/L +/-3.0, D/V -4.8) and .5 μl of AAV_5_-hSyn-DIO-rM3D(Gs)-mCherry (UNC Viral Vector Core) was infused at .1μl/min using a microsyringe pump. After infusion, syringes remained in place for 15 min before being slowly withdrawn. After bilateral infusion, incisions were sutured closed using nylon monofilament (Ethicon). For all surgeries, body temperature was maintained using a heating pad. After completion of surgery, anesthesia was reversed using Antisedan (atipamezole) and mice were allowed to recover on heating pads.

### Drug Administration

Clozapine-N-Oxide (Sigma) was diluted in sterile saline and administered at 1 mg/kg i.p. 30 min prior to behavioral testing. Istradefylline (Tocris # 5417) was dissolved in DMSO and diluted to 10% DMSO, 1% NP-40 in sterile saline immediately prior to i.p. administration at 3 mg/kg. Velneperit (MEdChem Exprpess #342577-38-2) has very low solubility in water, thus it was dissolved in pure DMSO prior to injection and injected at 100 mg/kg in .03 ml using an insulin syringe. Bexarotene (Tocris #5819) also has limited solubility in water, thus it was dissolved in pure DMSO prior to injection and injected i.p. at 50 mg/kg in a volume of .03 ml using an insulin syringe. These volumes of pure DMSO have been previously tested and validated to cause no adverse health effects in adult mice.

### Behavioral Assays

#### Auditory Cue-Dependent Fear Conditioning

Mice were habituated to fear conditioning chambers (Med Associates Inc., St Albans, VT) for 10 min each of two days prior to fear conditioning. Mice were conditioned to five tones (30s, 6 kHz, 65-70db) co-terminating with a 1 s foot shock (.65 mA, 1 mA for *Drd2* expression experiment, or .4 mA for mild conditioning).

#### Auditory Cue-Dependent Extinction

Cue-dependent fear extinction was tested 24 h after fear conditioning and extinction retention occurred 24 h after fear expression. For extinction, mice were placed in a novel context with a different olfactory cue, lighting and flooring and exposed to 15 or thirty 30 s, 65-70 db tones with an inter-trial-interval of 60 s. Freezing was measured using Freeze View software (Coulbourn Instruments Inc., Whitehall, PA).

#### Open Field

Open field chambers (Med Associates) were placed in a dimly lit room. Mice were placed in the chamber for 10 min and allowed to explore.

#### Brain collection following behavior for qPCR analysis of Drd2

Examination of changes in *Drd2* expression following behavioral experiments included 4 groups: 1) a Home Cage (HC)control group that remained undisturbed in their home cage throughout the experiment; 2) the primary experimental group (FC30), which received fear conditioning and extinction (30 CS) as described above; 3) a tone-alone control group (HC30) that remained in the home cage during training but was exposed to the same 30 tone presentations as the FC30 group in the extinction context; 4) a conditioned control group (FC1) that was fear conditioned as in the FC30 group but only exposed to one tone 24 h later. Brains were extracted 2 h after fear extinction or tone exposure. Brains from HC control animals were also extracted during this time.

#### Real Time PCR

RNA was reverse transcribed using SuperScript 4 (Invitrogen). Quantitative PCR was performed on cDNA with each sample run in triplicate technical replicates. Reactions contained 12 μl Taqman Gene Expression Master Mix (Applied Biosystems), 1 μl of forward and reverse primer, 1 μl of 5 ng/ul cDNA, and 6 μl water. Primers were proprietary FAM labeled probes from Life Technologies. Quantification of qPCR was performed on Applied Biosystems 7500 Real-Time PCR System. Cycling parameters were 10 min at 95°C, 40 cycles of amplification of 15 s at 95°C and 60 s at 60°C, and a dissociation step of 15 s 95°C, 60 s at 60°C, 15 s 95°C. Fold changes were calculated as ΔΔCT values normalized to levels of GAPDH or 18S mRNA. Values presented as fold change +/-s.e.m.

#### RNA-Seq Library Preparation

Libraries were generated from 1ng of Total RNA using the SMARTer HV kit (Clonetech), barcoding and sequencing primers were added using NexteraXT DNA kit. Libraries were validated by microelectrophoresis, quantified, pooled and clustered on Illumina TruSeq v3 flowcell. Clustered flowcell was sequenced on an Illumina HiSeq 1000 in 50-base paired end reactions. Approximately 25 million sequencing reads were collected per sample.

#### Analysis of RNA sequencing data

RNA sequencing data was analyzed using Tuxedo DESeq analysis software. Differential expression between HC and FC groups were obtained and used for further analysis. Using the q value of less than .05 as a cut-off, only highly significant returns were used for further analysis. To ensure that genes had a large enough difference in expression to warrant pharmacological manipulation, only those with differences in expression greater than 2^.5^ or ~141% were considered.

#### Bioinformatics

Enrichment analysis for Mouse Gene Atlas dataset and Jensen Compartments, was performed with Enrichr (Chen, E. Y. et al.).

Enrichment analysis for Gene Ontology Processes and Diseases was performed using MetaCore (Clarivate) Gene Set Enrichment Analysis was used to identify gene with concordant directional effects (Subramanian, A. et al.). Weighted gene network analysis was performed using GeneMania at the default setting (ref: Warde-Farley, D. et al.). Network data are presented in Dataset S3 A–C and were visualized in Cytoscape (ref: Montojo J, et al. (2010) GeneMANIA Cytoscape plugin: Fast gene function predictions on the desktop. Bioinformatics 26(22):2927–2928.) and presented in Fig. 4. Next using the MetaCore “Drugs for Drug targets” ‘Drug Gene Interaction Database’ (http://www.dgidb.org/) returns were examined for having a known pharmacological agent that modifies its activity. Genes lacking viable pharmacological modulators were eliminated.

#### Translating Ribosome Affinity Purification

TRAP procedure was completed as described in Heintz et. al., (2014)^30^. Adult Drd2-TRAP mice were anesthetized; their brains removed and snap frozen. Bilateral 1mm punches were collected and pooled from 3 animals per sample (n= 3 (HC) and 4 (FC)). Messenger RNA was isolated from eGFP-tagged ribosomes, as described in reference^30^. RNA was assessed for quality using the Bioanalyzer Pico (Agilent, Santa Clara, CA). All samples returned RINs (RNA Integrity Numbers) of 8.5 or greater.

#### Statistics

Statistical analyses were performed using Prism 6 or 7 by Graph Pad. All data presented as mean +/-s.e.m. Fear extinction experiments were examined using a repeated-measures ANOVA with drug as the between-subjects factor and tone presentation as the within subject factor. Open field activity or acoustic startle for Istrafefylline experiments was compared using a repeated measures ANOVA and a Tukey’s multiple comparisons analysis. For qPCR delta delta CT’s of data were compared by Student’s t-test between bound and unbound fractions. For all tests statistical significance was set at p < .05. For quantification of FISH RNA-Scope results, numbers of expressing verses co-expressing cells were compared using Mann-Whitney’s test.

#### FISH - RNA Scope Staining

Staining for RNA of interest was accomplished using RNA Scope Fluorescent Multiplex 2.5 labeling kit. Probes utilized for staining are: mm-Nts-C1, mmNts-C2, mm-Tac2-C2, mm-Sst-C1, mm-Sst-C2, mm-Crh-C1, mm, Prkcd-C1, mm-Prkcd-C3, mm-Drd2-C3, mm-Dkk3-C1, mm-Drd1a-C2, mm-Adora2a-C1. Brains were extracted and snap-frozen in methyl-butane on dry ice. Sections were taken at a width of 16 μm. Procedure was completed to manufacturers specifications.

#### Image acquisition

Images were acquired with the experimenter blinded to the probes used. 16-bit images of staining were acquired on a Leica SP8 confocal microscope using a 10x, 20x, or 40x objective. Images for 1A-F and 4B and E were acquired using a Zeiss Imager a1 with a 2x or 4x objective. Within a sample, images used for quantification were acquired with identical settings for laser power, detector gain, and amplifier offset. Images were acquired as a z-stack of 10 steps of .5 μm each. Max intensity projections were then created and analyzed.

## Acknowledgements

Support was provided by NIH (R01 MH108665-01) and Cohen Veteran Biosciences foundation.

---

**Supplemental Figure 1.**
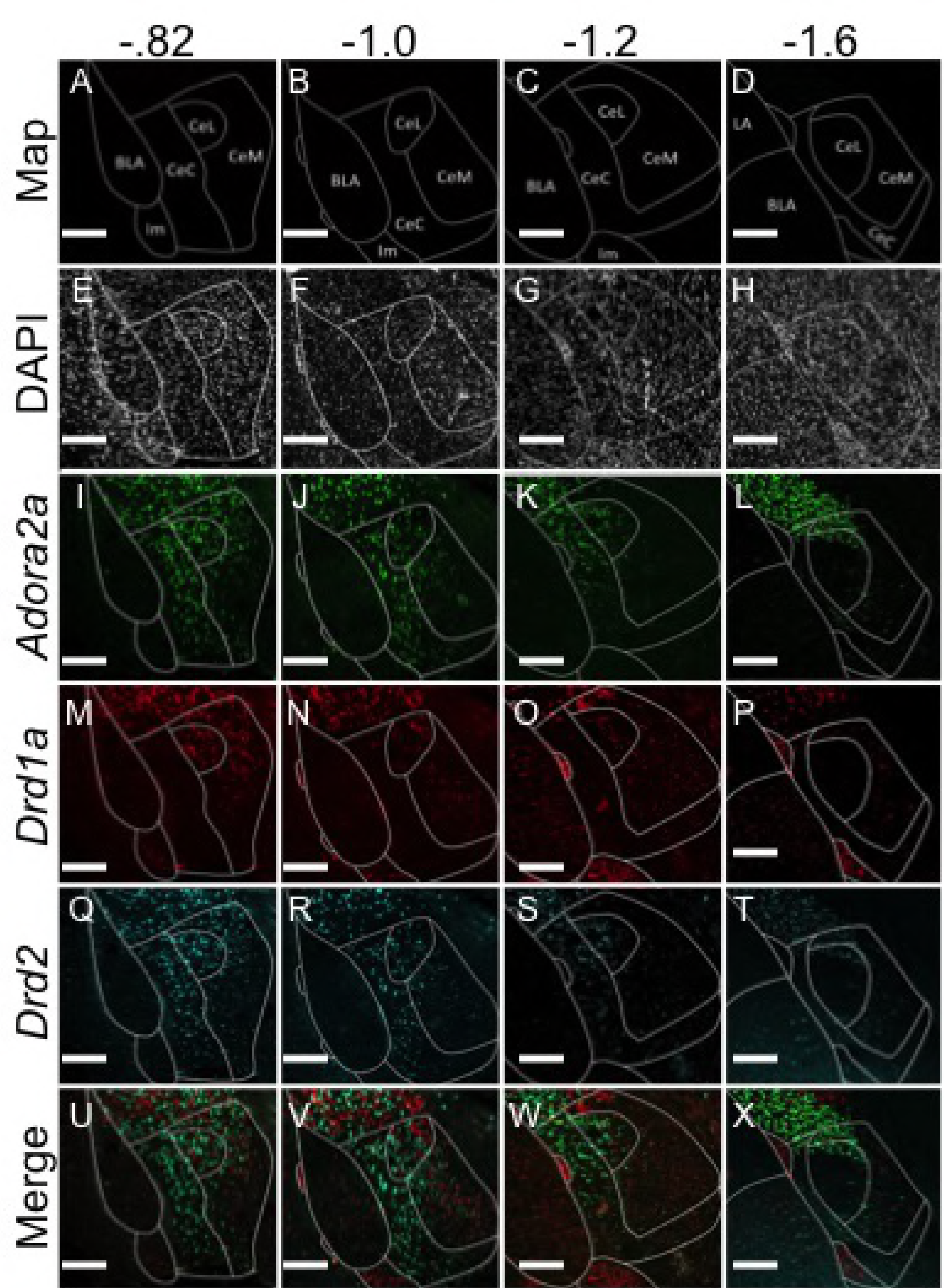
Distribution of *Drd2, Adora2*, and *Drd1a* across A/P axis of CeA. The *Drd2/Adora2a* population is heavily represented in anterior CeC and CeL, while *Drd1a* is strongly represented in anterior CeL and CeM. Neither population is found at high levels in posterior CeA. **A-D**. Map of CeA at A/P: -.82, -1.0, -1.2, and -1.6 respectively. **E-H**. DAPI (Grey) expression at A/P: -.82, -1.0, -1.2, and -1.6. **I-L**. *Adora2a* (Green)expression at A/P: -.82, -1.0, -1.2, and -1.6. *Adora2a* is strongly expressed in anterior CeC and CeL (-8.2 to -1.2) but not in posterior CeA (-1.6). **M-P**. *Drd1a* (Red)expression at A/P: -.82, -1.0, -1.2, and -1.6. *Drd1a* is strongly expressed in anterior CeL and CeM (-8.2 to -1.2) but not in posterior CeA (-1.6). **Q-T**. *Drd2* (Cyan) expression at A/P: -.82, -1.0, -1.2, and -1.6. *Drd2* is strongly expressed in anterior CeC and CeL (-8.2 to -1.2) but not in posterior CeA (-1.6). **U-X**. Merge of channels. *Drd2* and *Adora2a* form a single co-expressing population found primarily in anterior CeC and CeL that does not co-express with *Drd1a*, which is found primarily in anterior CeL and CeM. Scale Bar: A-X 200 um.

**Supplemental Figure 2.**
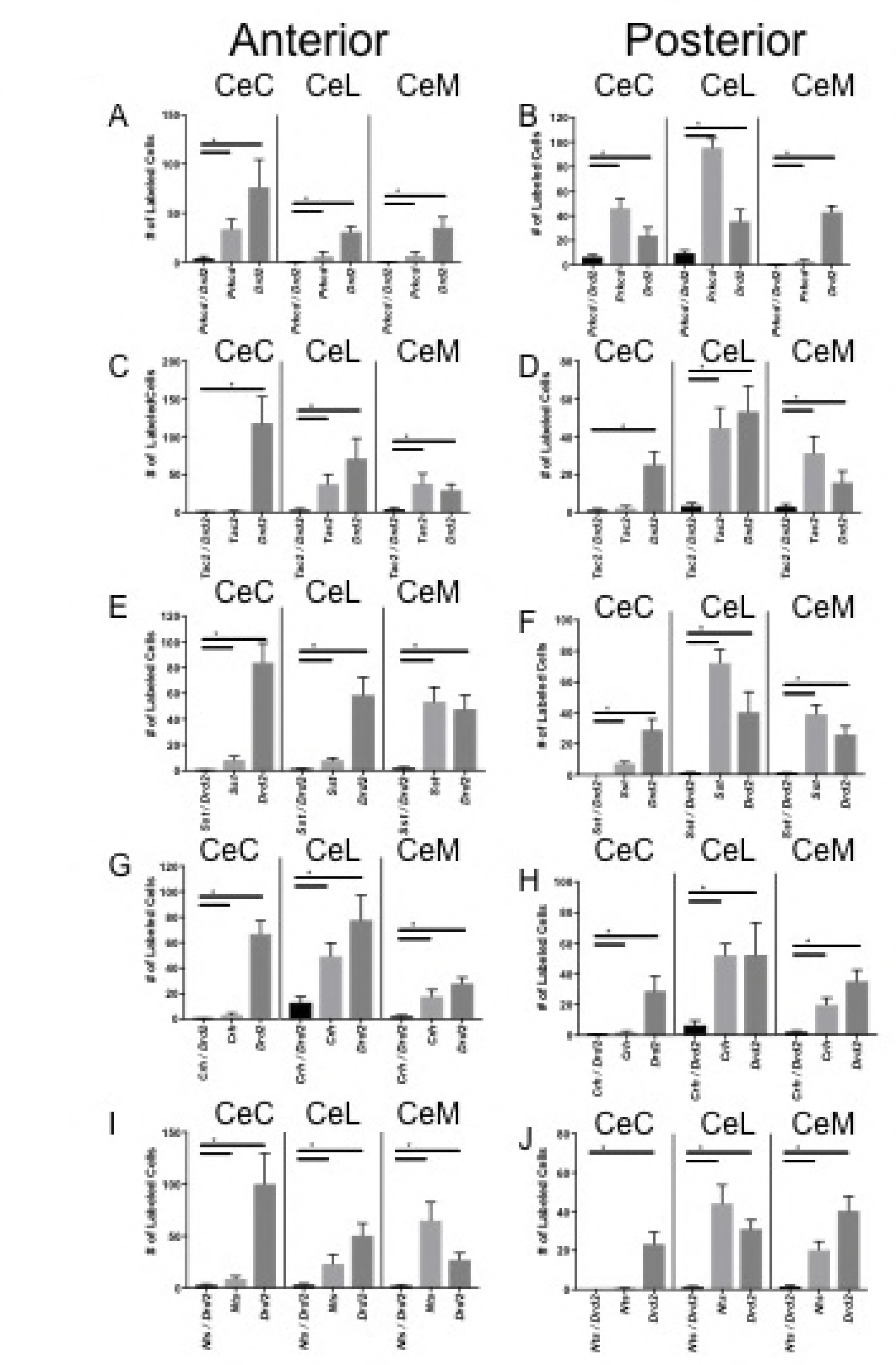
Quantification of Co-expression of *Drd2* with *Prkcd, Tac2, Sst, Crh*, and *Nts* in anterior and posterior CeA. *Drd2* co-expression was quantified and represented as total number of cells expressing *Drd2*, expressing other RNA of interest, or co-expressing the two RNA’s. In all cases Mann-Whintney test was performed to determine whether the co-expressing population was significantly different from each single expressing population. * p<.05. **A**. In all regions of anterior CeA very little co-expression between *Drd2* and *Prkcd* was found. **B**. Moderate co-expressoin between *Drd2* and *Prkcd* was found in posterior CeC and CeL. **C-J**. Very limited coexpression was detected between *Drd2* and *Tac2, Sst, Crh*, and *Nts*, and in all cases the *Drd2* labeled population is significantly different from the co-labeled population.

**Supplemental Figure 3.**
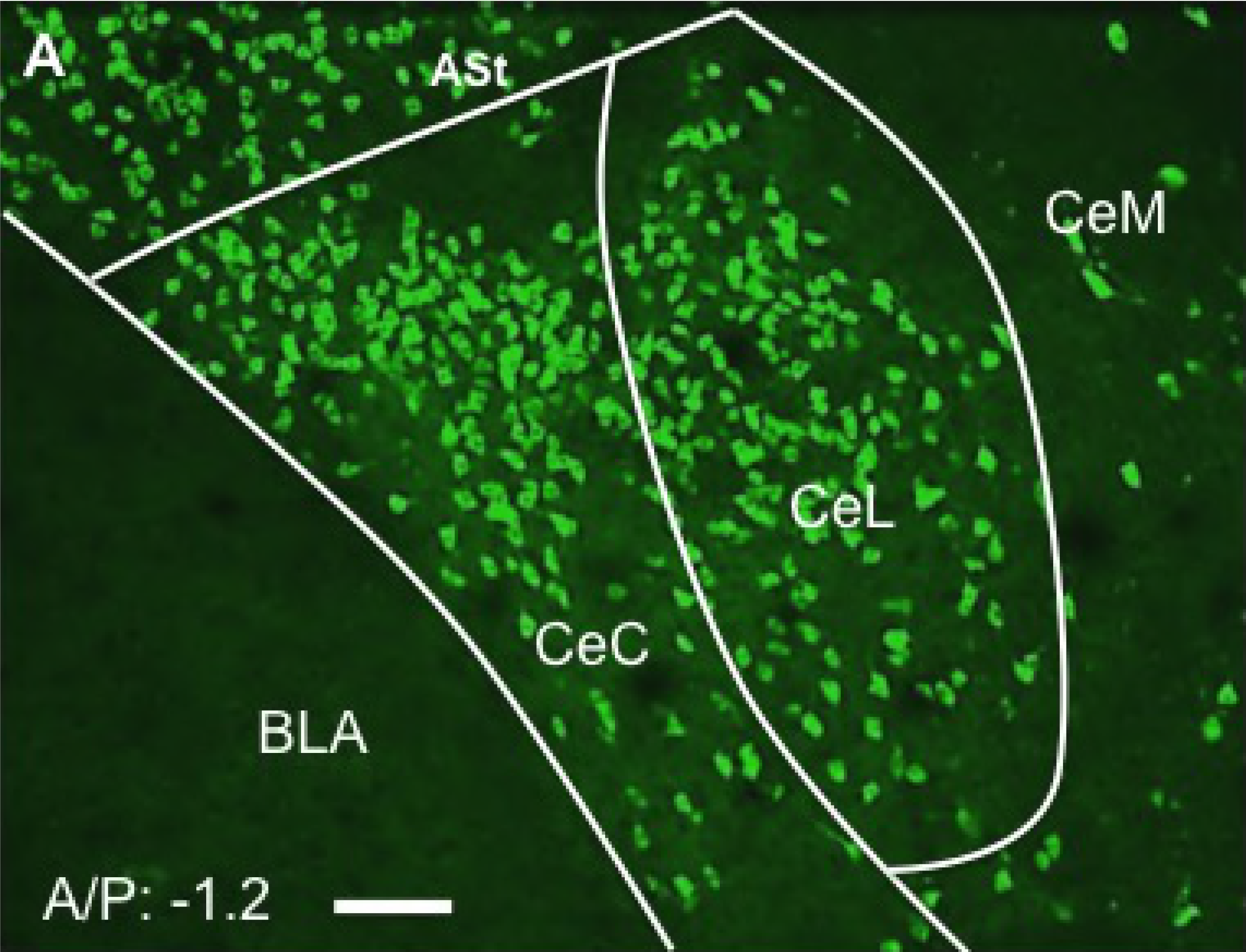
Drd2-TRAP transgene expression closely recapitulates Drd2expression pattern observed with FISH. **A**. L10a-GFP ribosomal subunit (Green) is expressed is pattern very similar to that observed with FISH.

**Supplemental Figure 4.**
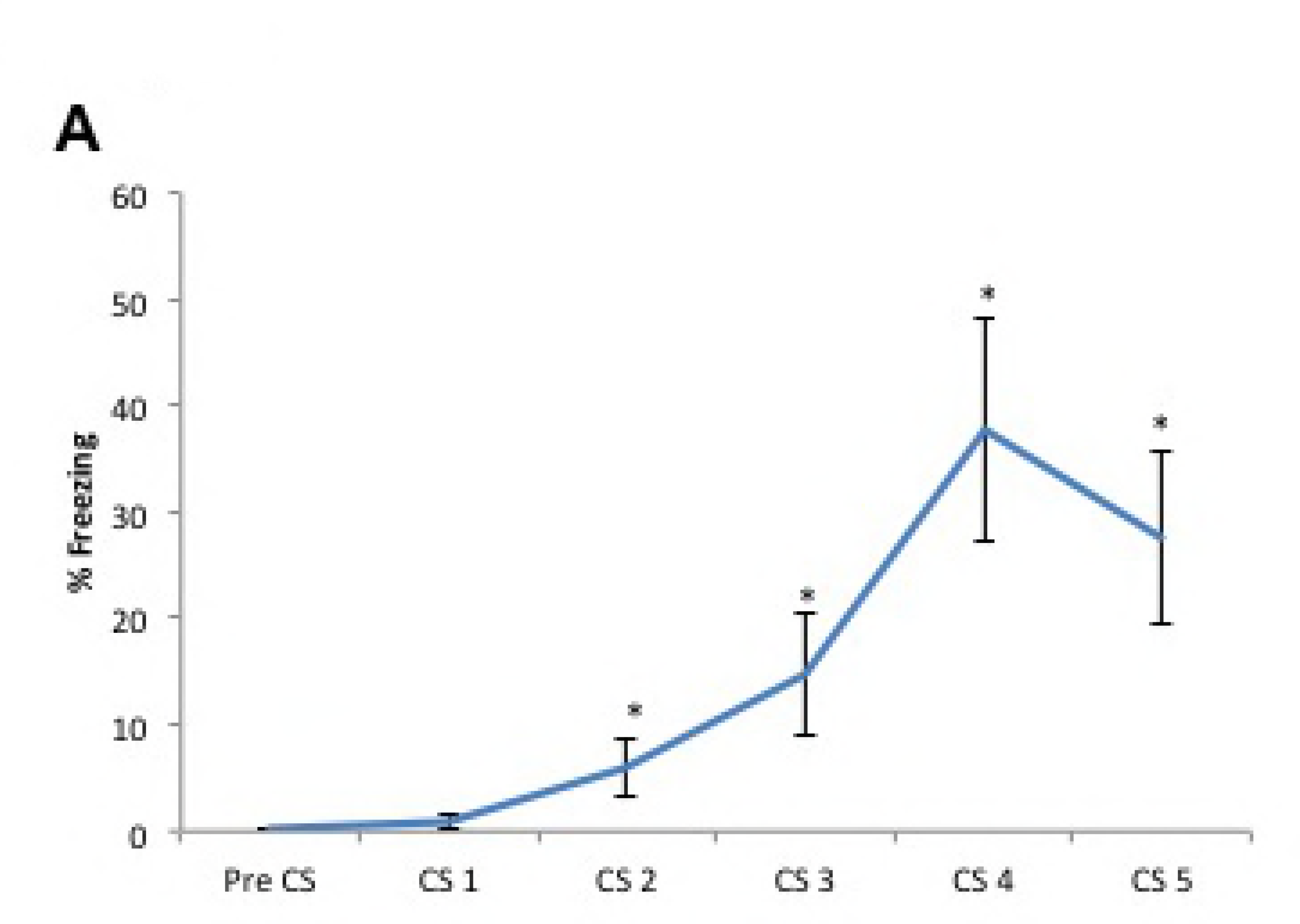
Fear conditioning of mice for TRAP collection. Mice express significantly more freezing after fear conditioning. (Unpaired t-Test, freezing to tone vs. CS1, * p < 05).

**Supplemental Figure 5.**
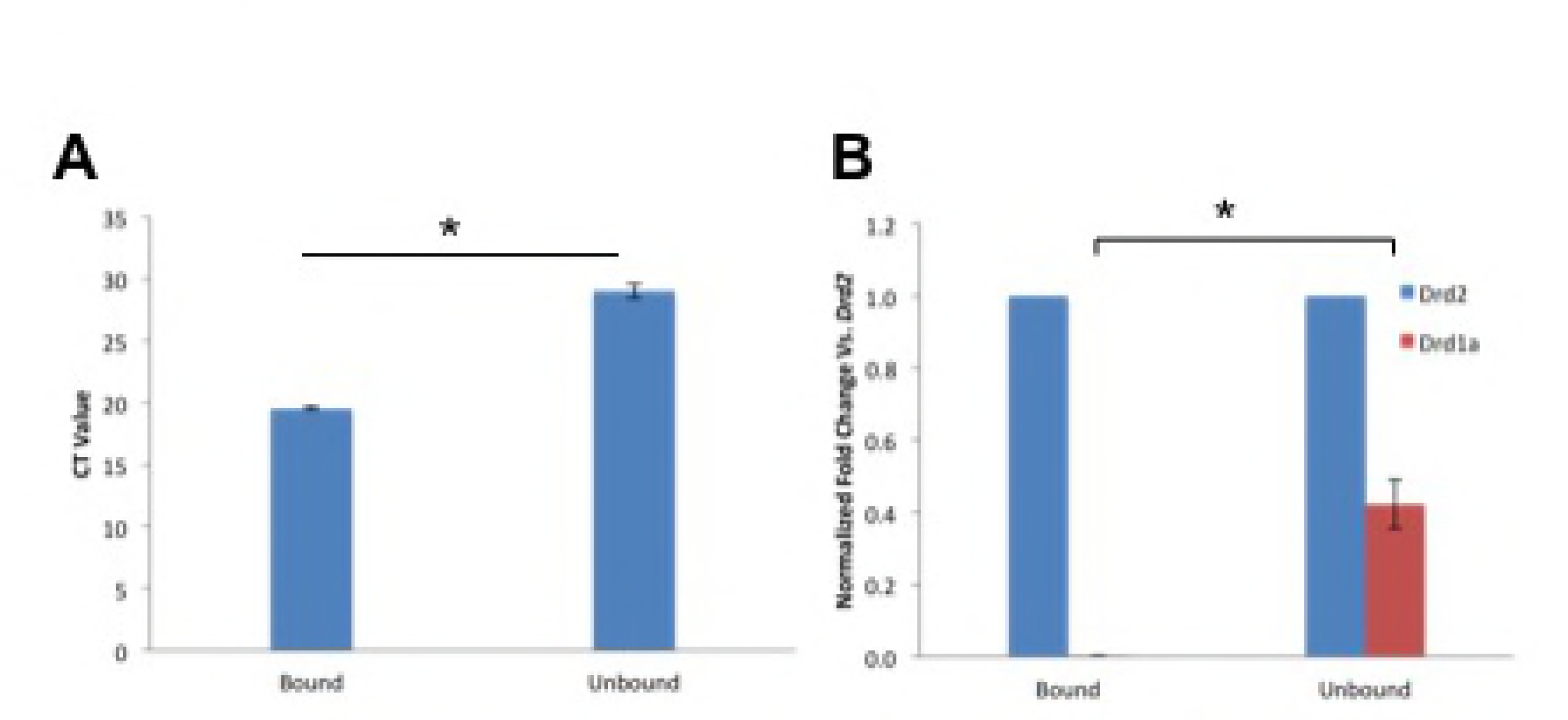
Validation of TRAP pull-down. **A**. Ribosomal subunit 18S is found at significantly higher levels in bound fraction, verifying ribosomal pull down (Paired t-Test, * p < .05). **B**. The ratio of *Drd2:Drd1a* is significantly higher in bound fraction vs. unbound fraction, verifying RNAs were successfully isolated from *Drd2* neurons (Paired t-Test, * p < .05).

**Supplemental Figure 6.**
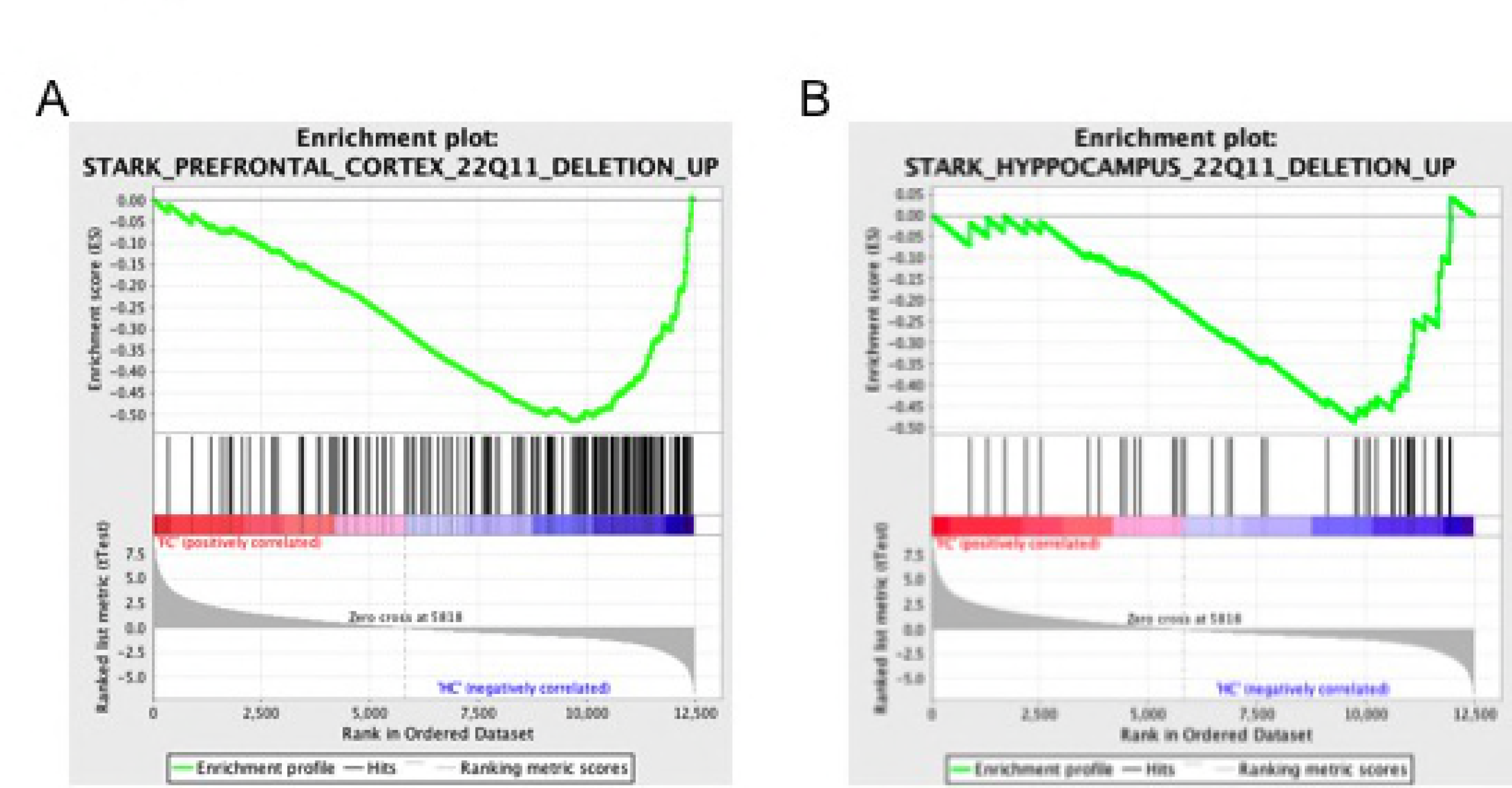
Gene set enrichment analysis. Using the entire expression dataset in default setting genes identified in *Drd2*-TRAP fear conditioning study and humanized model of 22q11.2 deletion are significantly and concordantly regulated in inverse direction in both **A**. Prefrontal cortex and **B**. Hippocampus.

**Supplemental Figure 7.**
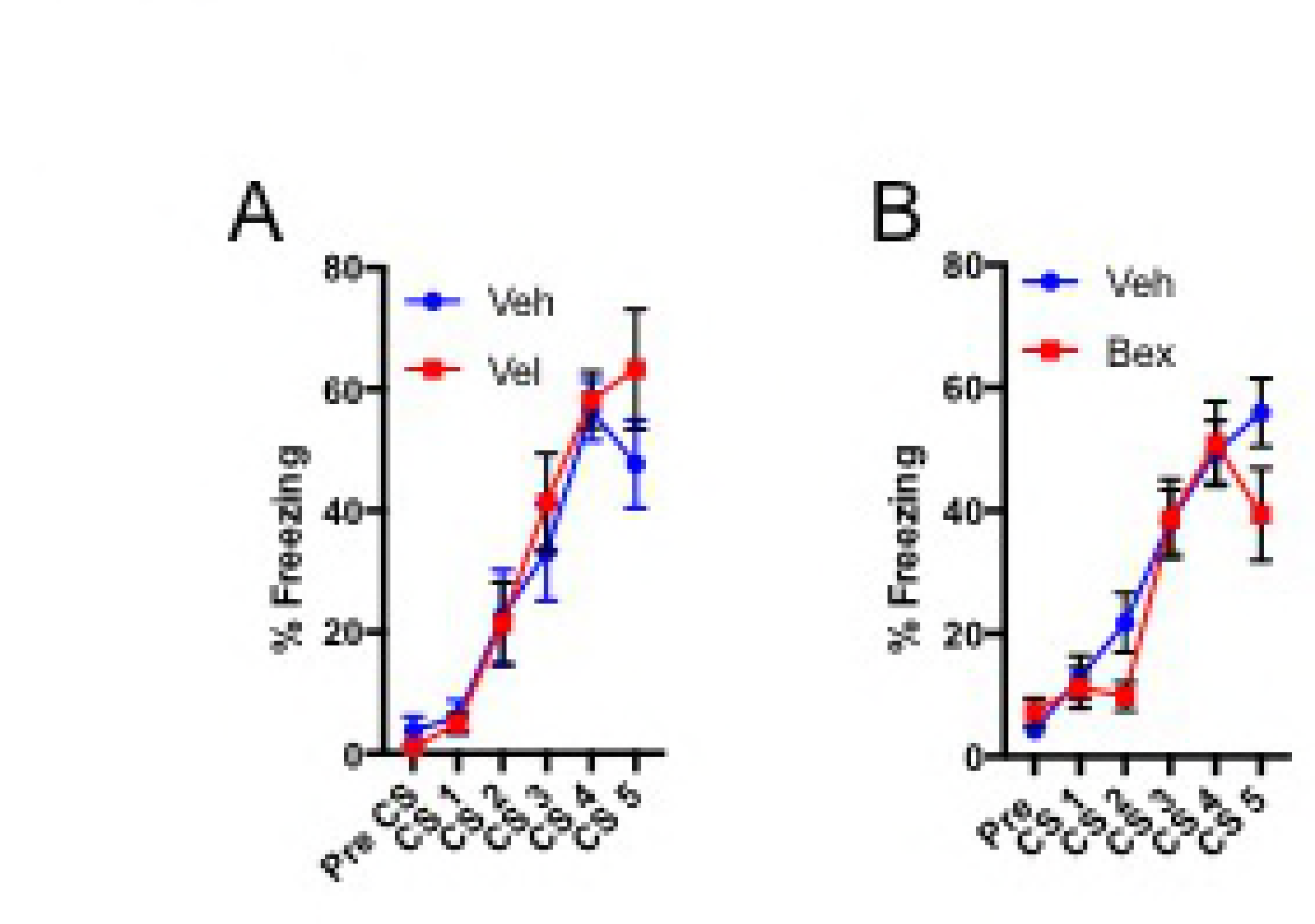
Fear conditioning for Veleneperet and Bexarotene experiments. In each case two groups of animals were fear conditioned (5 CS/US, .65 mA). **A**. Animals later given Velenperit were fear conditioned. No significant differences between groups were detected(2-way RM ANOVA, F(1,13)=.3773, p > .05). **B**. Animals later given Bexarotene were fear conditioned. No significant differences between groups were detected(2-way RM ANOVA, F(1,30)=.712, p > .05).

**Supplemental Figure 8.**
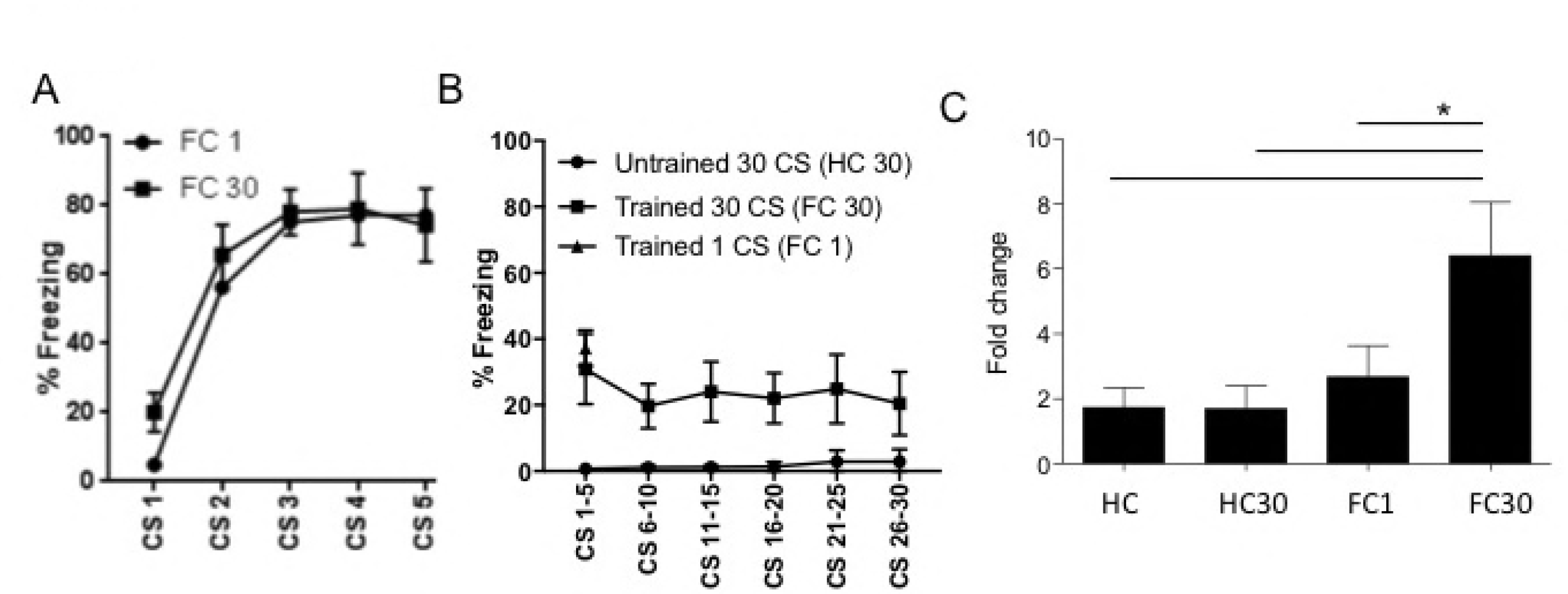
Examination of *Drd2* expression following fear conditioning and extinction. **A**. Two groups of animals (FC1 and FC 30) were fear conditioned (5 CS/US, .65 mA US). **B**. Twenty-four hours three groups of animals were exposed to extinction context. FC1 was exposed to a single CS in extinction context. Percent freezing indicates freezing during tone during CS 1 and freezing during the corresponding time when tone was played for other groups for CS 2-30, freezing during CS 1 was significantly greater than control (unpaired t-Test F(1,12)=5.929, p <.0001). HC 30 group did not receive fear conditioning, but was exposed to 30 CS’s in extinction context. FC 30 was exposed to 30 CS’s in extinction context and expressed more freezing than HC 30 group (Each bin examined by students t-test p <.05). **C**. Four groups were sacrificed 2 hours following behavior and qPCR performed on amygdala punches. FC30 group had significantly increased *Drd2* expression compared to all other groups.

**Supplemental Table 1.**
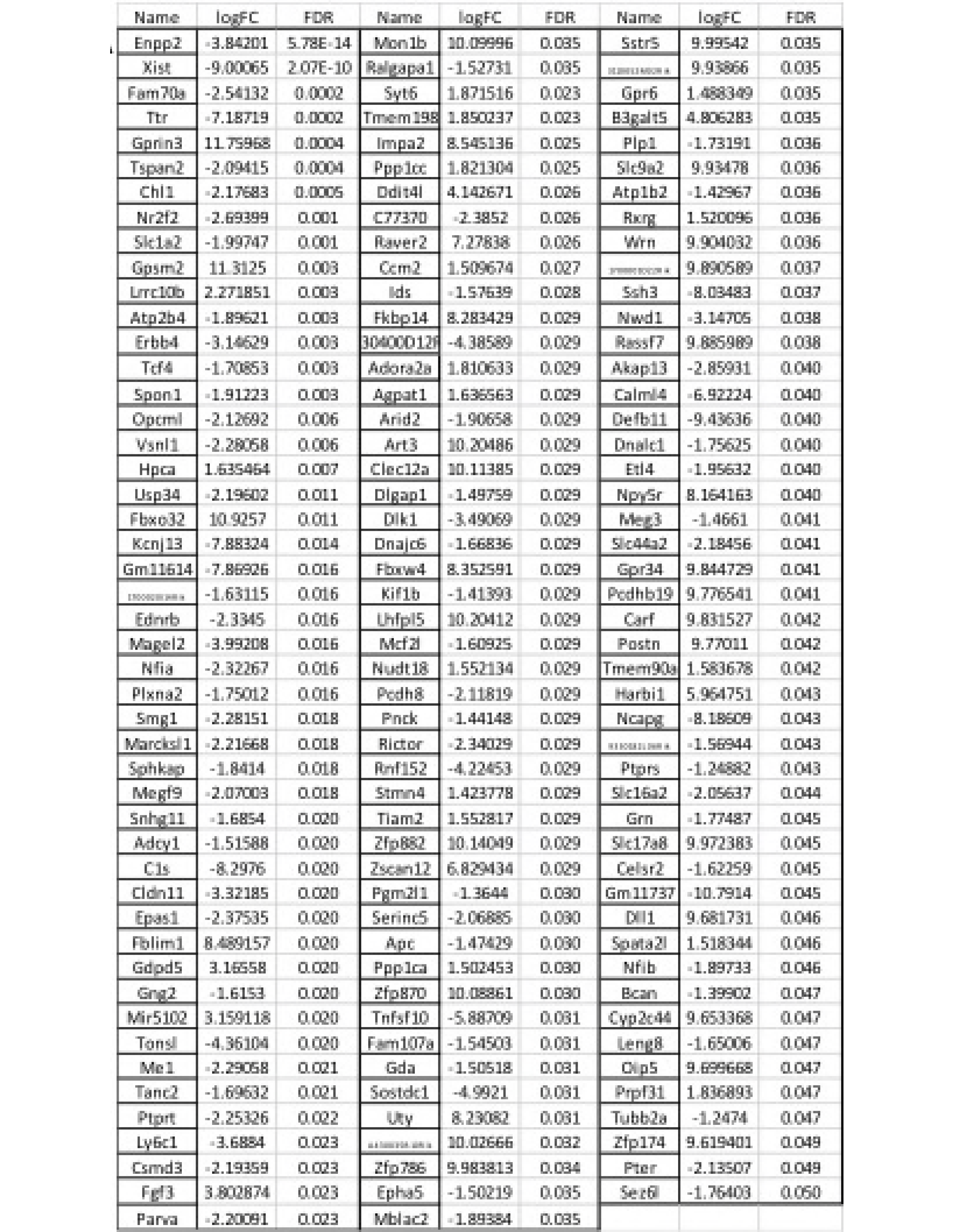
Complete list of differentially regulated RNA’s. **A**. LogFC indicates log fold change between FC and Control group. Negative LogFC’s indicate decreased expression compared to control while positive values indicate increased expression. FDR indicates false discovery rate corrected across multiple tests. Only q< .05 values are featured.

**Supplemental Table 2.**
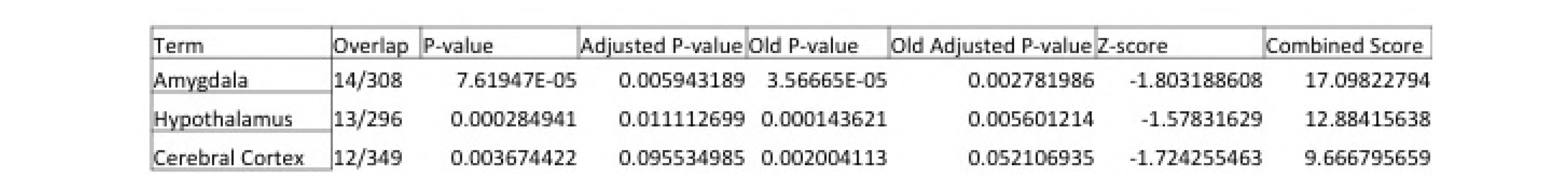
Using Mouse Gene Atlas dataset, Enrichr confirms amygdala specificity of pull-down and gene change.

**Supplemental Table 3.**
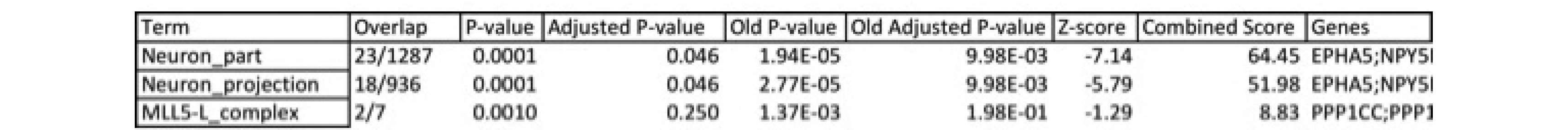
Jensen COMPARTMENTS analysis. Jensen COMPARTMENTS analysis data set, using sequence based prediction methods confirms neuronal specificity of pull-down and gene change.

**Supplemental Table 4.**
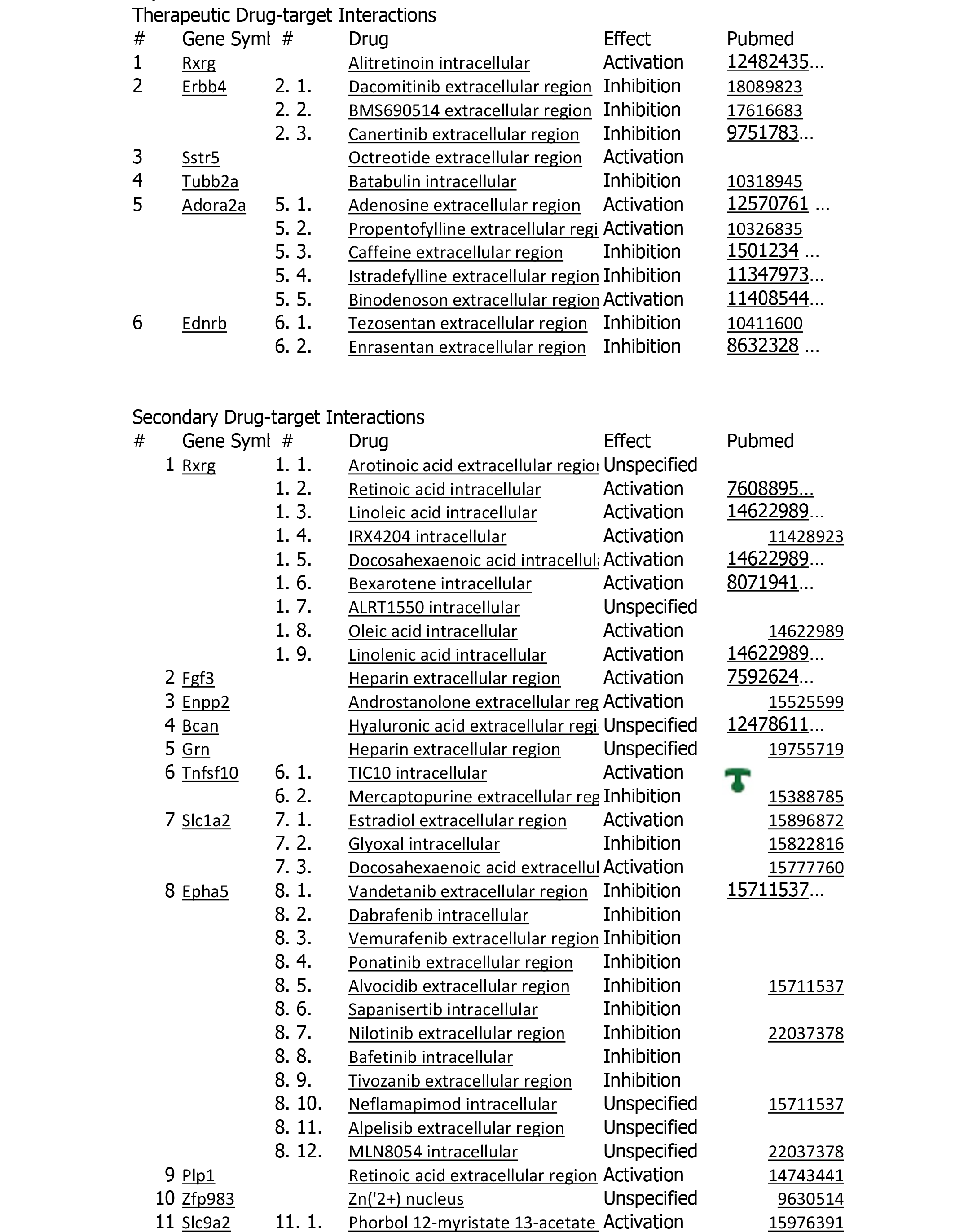
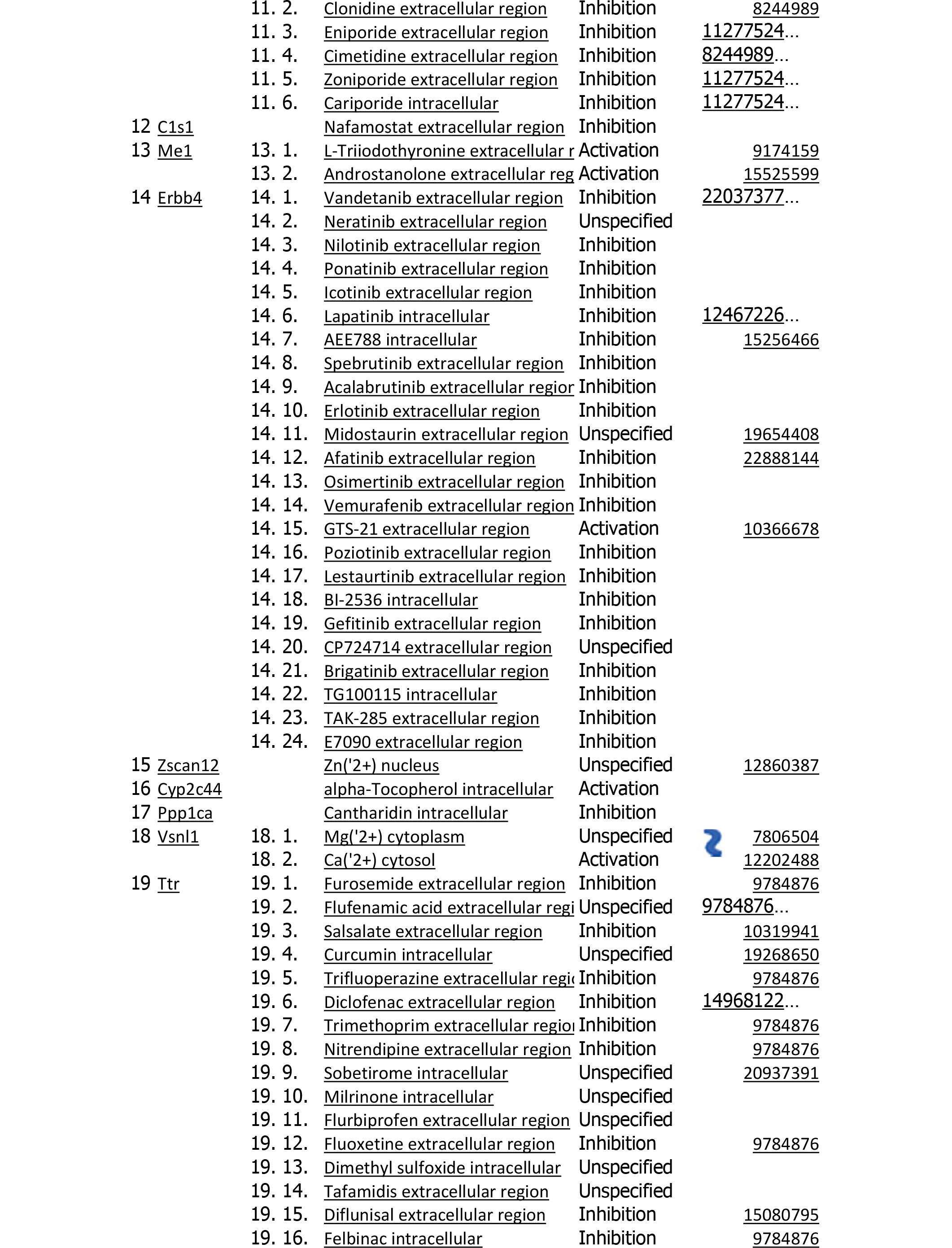
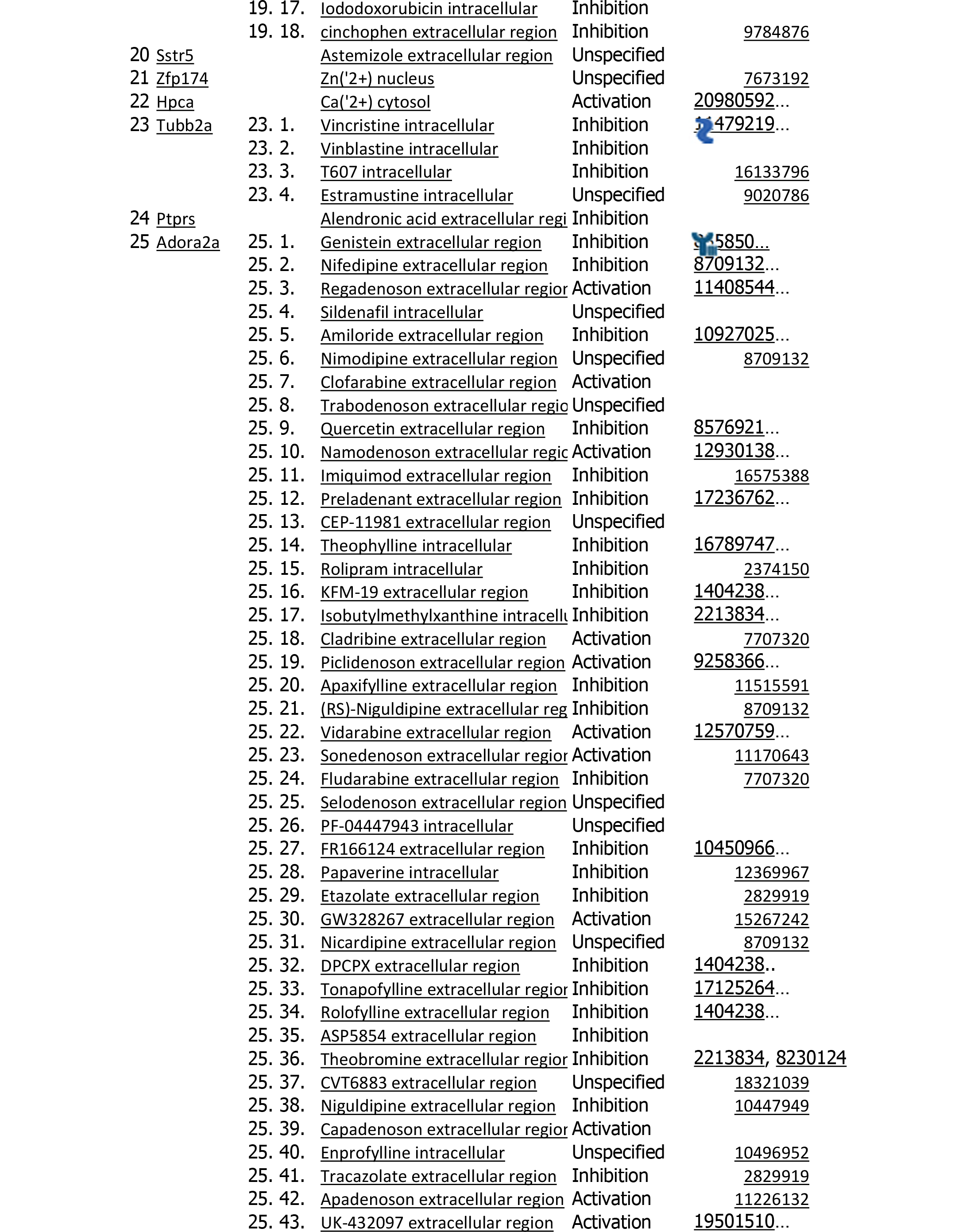
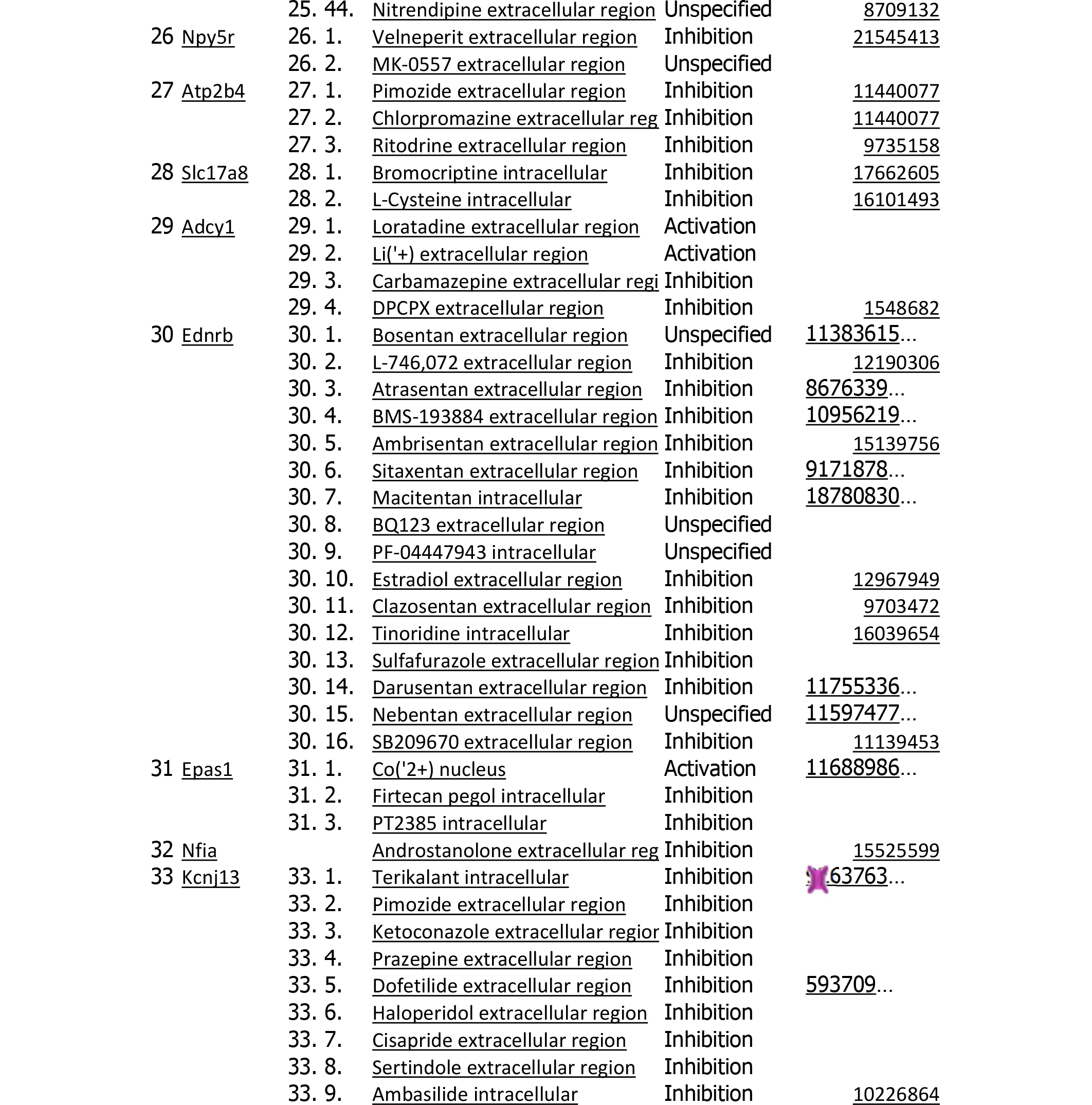
Drug-Gene analysis of differentially expressed genes. Metacore ‘Drug for Drug database’ identifies drugs that target protein products of differentially expressed genes.

